# Consequences of intra-locus recombination for branch-length-based inference of gene flow

**DOI:** 10.64898/2026.08.10.743750

**Authors:** Arthur Boddaert, Bert Van Bocxlaer, Camille Roux

## Abstract

Phylogenomic methods provide a powerful way to study introgression across broad clades of the tree of life, because they can test for gene flow from gene trees without requiring population-level resequencing data. These methods generally assume that each locus can be represented by a single non-recombining genealogy, which may be violated when recombination occurs within loci. Here, we used coalescent simulations to evaluate how intra-locus recombination affects gene-flow inferences in `Aphid`, a method using branch lengths to distinguish gene flow from incomplete lineage sorting in species triplets. Across the conditions tested, `Aphid` accurately recovered the proportion of loci affected by recent and intermediate gene flow, while recombination reduced the underestimation observed when gene flow is ancient. It also retained a relative timing signal, with accuracy decreasing as gene flow became older. This relative-timing approach was then applied to 456 African cichlid exon trees, where proposed gene flow involving *Coptodon* was consistently associated with intermediate-to-old rather than recent gene flow. Overall, our simulations suggest that intra-locus recombination does not increase error in `Aphid`’s inference of the prevalence of gene flow under the conditions tested, but can reduce temporal resolution for intermediate and ancestral events. When applied to cichlids, we show that this loss of resolution still permits the distinction between recent and older gene-flow.

## Introduction

Large-scale genomic datasets are deeply changing the way species histories are reconstructed. Instead of providing a single, uniform picture of divergence, genomes often reveal a mosaic of phylogenetic histories, with different chromosomes, or even different regions within chromosomes, supporting distinct genealogical relationships (Boussau and Scornavacca, 2020). Such heterogeneity has been documented at broad genomic scales, for instance between autosomes and sex chromosomes (Patterson et al., 2006), but also at finer scale, where recombination, drift, selection, and introgression jointly shape local genealogies (Roux et al., 2013). As a consequence, conflicts between gene trees and the species tree have become a central topic in phylogenomics (Pease et al., 2016; Malinsky et al., 2018; Edelman et al., 2019; Glémin et al., 2019; Vanderpool et al., 2020). Gene flow (GF) and incomplete lineage sorting (ILS) are the two major biological processes causing such conflicts. GF transfers genetic material between diverging lineages and can cause gene trees to depart from the species history (Doyle, 1992). ILS, by contrast, occurs when genetic lineages do not coalesce in the ancestral population immediately preceding a speciation event but instead coalesce deeper in the species tree (Maddison, 1997). Distinguishing between these two processes is therefore essential for interpreting discordance among gene trees and for reconstructing the species history accurately.

A widely used strategy to distinguish GF from ILS relies on the expected symmetry of discordant topologies in gene trees under ILS alone. For a species tree, ((*A, B*), *C*), ILS, in the absence of gene flow, is expected to generate the two alternative discordant topologies ((*A, C*), *B*) and ((*B, C*), *A*) at equal frequencies. An excess of one discordant topology over the other has therefore been widely used as evidence for introgression. This rationale underlies ABBA-BABA tests and the *D* statistic, which test for introgression from asymmetries in allele-sharing patterns across species quartets (Green et al., 2010; Durand et al., 2011), as well as several extensions of this framework (Martin et al., 2015). These approaches are powerful, but they mainly rely on topology or site-pattern asymmetry. They become uninformative when gene flow produces balanced topological discordance, for instance when introgression occurs symmetrically or when multiple migration events generate opposite signals (Vanderpool et al., 2020). Under such scenarios, branch lengths provide an additional source of information, because they contain information about coalescent times and can help to distinguish the contribution of GF and ILS in generating discordant topologies, regardless of the level of symmetry in GF.

Two methods that use branch-length information from discordant gene trees to distinguish GF from ILS have recently been developed: `QUIBL` (Edelman et al., 2019) and `Aphid` (Galtier, 2024). Both methods exploit the expectation that discordant gene trees generated by ILS and GF differ in their branch-length distributions. Under ILS, gene-tree discordance arises because lineages fail to coalesce in their most recent ancestral population and instead persist until a deeper ancestral population, where the first coalescence eventually occurs. The resulting delay produces longer terminal branches and a greater total tree length. By contrast, under GF, discordance can result from the transfer of genetic material between non-sister lineages, bringing them into the same population more recently than expected under a strictly allopatric model of divergence. GF-associated discordant trees are therefore expected to be shorter than discordant trees generated by ILS, especially in their terminal branches. Whereas `QUIBL` and `Aphid` both use branch-length distributions to estimate the relative contribution of GF and ILS, they differ in their analyzed source of information. `QUIBL` mainly examines the distribution of internal branch lengths, whereas `Aphid` also uses variation in terminal branches. This makes `Aphid` particularly relevant for evaluating not only the amount of GF inferred from discordant gene trees, but also whether branch-length information retains a signal about the relative timing and direction of GF. A central assumption of `Aphid`, and other gene-tree-based phylogenomic methods including `QUIBL`, is that each analysed locus can be represented by a single genealogy (Galtier, 2024). This assumption is common to many coalescent-based phylogenetic methods, in which each locus is treated as an independent genealogical unit (Heled and Drummond, 2010; Wen and Nakhleh, 2018; Edelman et al., 2019). In practice, this assumption implies that intra-locus recombination is absent, or sufficiently weak to be ignored at the scale of the analysed loci. Recombination can generate a mosaic of local genealogies along a genomic sequence, however, so that different positions within the same locus may have different ancestries (Hudson, 1983; Hudson and Kaplan, 1985; Posada and Crandall, 2002; Wakeley, 2020). Under these conditions, the tree inferred for a recombining locus may no longer correspond to a single genealogy, but reflect a composite of several local histories (Griffiths and Marjoram, 1996; McVean and Cardin, 2005; Wakeley, 2020).

In coalescent theory, the ancestry of a recombining locus is formally described by the ancestral recombination graph (ARG), from which a local genealogy can be extracted at each position along the sequence. In this framework, recombination mainly affects the dependency among neighbouring genealogies: positions that are close to each other tend to share similar histories, but this tendency decreases as the recombination distance between sites increases (Wiuf and Hein, 1999; Hudson and Kaplan, 1985). However, recombination does not change the marginal distribution of the local genealogy at a given position. The topology and branch lengths of a local tree considered in isolation therefore follow the same distribution as those of a non-recombining locus evolving under the same demographic model (Griffiths, 1999; Wakeley, 2020). Therefore, coalescent theory suggests that, if `Aphid` were applied to true local genealogies, recombination alone would not be expected to generate a systematic bias in GF inference. Practically, the question remains open, however, because gene-tree-based phylogenomic methods to infer GF like `Aphid` are not applied to true local trees, but to a single tree reconstructed for each locus. Whether this theoretical robustness carries over to reconstructed locus trees therefore remains unclear.

To assess whether this practical approximation matters, we simulated recombining loci for which the underlying local genealogies were known, but then represented each locus by a single tree, as required by `Aphid`. This design mimics the situation faced in empirical phylogenomic studies, where a locus affected by recombination is usually analysed through one inferred gene tree. It allowed us to ask whether intralocus recombination alters the branch-length signal used to distinguish GF from ILS, or whether the dominant genealogical signal of a locus remains sufficient for reliable inference. We evaluated three quantities derived from `Aphid`: the inferred proportion of loci affected by GF, a relative timing estimate developed here, and support for the direction of GF.

Although `Aphid` allows lineages brought together by GF to coalesce at different times, Galtier (2024) used this temporal component mainly to quantify the contribution of ancient scenarios. Here, we used the support assigned to these different times to distinguish recent from older GF-associated coalescence, and tested how recombination affected this distinction. We then applied the same approach to a published case of reticulation in African cichlids. Phylogenetic network analyses by Astudillo-Clavijo et al. (2023) inferred gene flow involving *Coptodon*, but did not determine whether the corresponding exchange was recent or occurred deeper in the history of these lineages. We therefore used this demographical context to ask which of these two temporal interpretations was better supported by the gene-tree branch lengths, and to illustrate how phylogenomic data can be used to distinguish recent from older GF.

## Materials and Methods

`Aphid` infers GF from observed phylogenetic conflict in species triplets by examining topological discordance between gene trees and the species tree, and the branch-length variation associated with this discordance (Galtier, 2024). It estimates the relative contribution of ILS and GF to the observed phylogenetic conflict. In addition, its scenario-based output can be used to extract information about the direction of GF and the relative timing of the coalescence events associated with GF. We used simulations to evaluate the accuracy of these inferences, focusing on the inferred proportion of loci affected by GF, the direction of GF, and its relative timing. We then tested how these inferences were affected by the amount of ILS and by intra-locus recombination.

### General Framework

We used coalescent simulations to evaluate how intra-locus recombination affects the interpretation of `Aphid` inferences. The objective was not to reproduce the full complexity of empirical phylogenomic datasets, but to generate controlled datasets in which the true history of each locus was known. This design allowed us to compare simulated and inferred histories while varying the main factors expected to influence `Aphid` inference: the proportion of loci affected by GF, the relative timing and direction of GF, the amount of ILS, and the intra-locus recombination rate.

All simulations were performed with the coalescent simulator `ms` (Hudson, 2002). Each simulated dataset contained 1,000 loci, and each parameter combination was replicated 100 times. We considered a focal species triplet with species tree ((*A, B), C*), for which GF was simulated between the non-sister lineages *A* and *C*. This triplet was embedded in a larger nine-species phylogeny, including (Fig. 1):

**Fig. 1.**
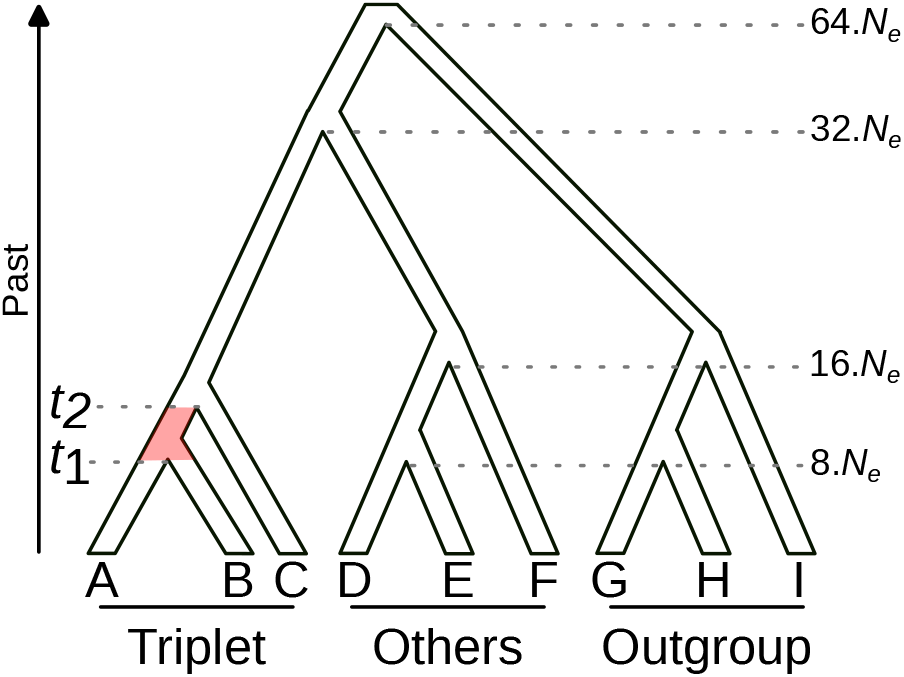
Species trees used to generate three levels of incomplete lineage sorting (ILS). The focal triplet is *((A, B), C)*, embedded in a larger nine-species phylogeny including three additional non-focal species and three outgroups. The amount of ILS was controlled by modifying the length of the internal branch (in red) between the *A*–*B* split (*t*_*1*_ = 8*N*_*e*_ generations) and the deeper split between (*A, B*) and *C* (*t*_2_). This interval was set to 8*N*_*e*_, 5*N*_*e*_, and 2*N*_*e*_ generations, corresponding to expected ILS probabilities of approximately 2%, 8% and 37%, respectively

1. the focal triplet used for GF inference,
2. three additional non-focal species, and
3. three outgroup species.

These additional species provided the phylogenetic information required for rooting gene trees and estimating locus-specific branch-length scaling factors.

For each GF scenario, simulations were repeated under three levels of ILS and three intra-locus recombination rates. The amount of ILS was controlled by changing the length of the internal branch separating the *A*–*B* split, denoted *t*_1_, from the deeper split between (*A, B*) and *C*, denoted *t*_2_ (Fig. 1). Recombination was then introduced within loci to generate several local genealogies along the same simulated sequence. Because `Aphid` analyses one tree per locus, these local genealogies were summarized into a single reconstructed locus tree before running `Aphid`. This step is central to our study, as it mimics the practical situation in which a recombining locus is represented by a single inferred gene tree, while ensuring that recombining and non-recombining simulations differ only by the presence of recombination, not by an additional phylogenetic reconstruction step that could introduce methodological biases.

We assumed an effective population size *N*_*e*_ = 10, 000 individuals. Trees simulated by `ms` have branch lengths expressed in coalescent units, whereas `Aphid` requires branch lengths expressed as numbers of substitutions per site. We therefore rescaled all branch lengths by multiplying them by 4 *N*_*e*_*µ*, assuming a mutation rate *µ*=10^−8^ per site per generation. Under these parameter values, all branch lengths were multiplied by 0.0004 before being analysed with `Aphid`.

### Species Tree and Levels of Incomplete Lineage Sorting

To test how the amount of ILS affects `Aphid` inferences, we simulated datasets under three levels of ILS. These levels were obtained by modifying the length of the internal branch of the focal species tree, that is, the time interval between the *A*–*B* split, denoted *t*_1_, and the deeper split between (*A, B*) and C, denoted *t*_2_. Under the multispecies coalescent, ILS occurs when the *A* and *B* lineages fail to coalesce during this interval and instead persist into the deeper ancestral population. The probability of this event is given by 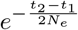. We therefore considered three internal branch lengths: 8*N*_*e*_, 5*N*_*e*_, and 2*N*_*e*_ generations. These correspond to expected ILS probabilities of approximately 2%, 8%, and 37%, respectively, in the absence of GF (Fig. 1). The three values span low to high ILS, from conditions in which most loci follow the species tree to conditions in which a substantial fraction coalesce deeper in the tree.

### Gene Flow Scenarios

Gene flow was simulated between the two non-sister lineages *A* and *C* of the focal triplet ((*A, B*), *C*). We considered four proportions of loci affected by GF: 0, 10, 25, and 50%. These values correspond to the proportion of loci for which a GF event was imposed in the simulation. For loci affected by GF, all events within a simulated dataset were assigned to the same relative time window. We considered three timing categories, expressed relative to *t*_1_, the time of the *A*–*B* split: recent GF, with reference time *t*_*GF*_ = 0.1*t*_1_; intermediate GF, with *t*_*GF*_ = 0.5*t*_1_; and ancestral GF, with *t*_*GF*_ = 0.9*t*_1_. Each category was implemented as a migration pulse over the corresponding time interval detailed in the Supplementary Methods. These categories were designed to test whether the timing information extracted from `Aphid` outputs can recover a relative temporal signal, rather than to estimate an absolute date of introgression.

GF was simulated as unidirectional. We analysed both directions between *A* and *C* separately, corresponding to *A* → *C* and *C* → *A*. These two scenarios generate the same class of discordant topology, ((*A, C*), *B*), but differ in the expected distribution of terminal branch lengths. Including both directions therefore allowed us to test whether intra-locus recombination alters the branch-length signal on which directional inference in `Aphid` would rely.

### Intra-Locus Recombination and Construction of Locus Trees

Like many gene-tree-based approaches, `Aphid` assumes that each analysed locus can be represented by a single non-recombining genealogy. To test how intra-locus recombination affects `Aphid` inferences, all simulations were performed under two recombination rates, in addition to a control without recombination. The population-scaled recombination parameter for the entire locus was defined as *ρ*=4*N*_*e*_*rL*, where *r* is the recombination rate per site per generation and *L* is the locus length in nucleotides. Consequently, the amount of within-locus recombination examined here reflects the combined effects of *N*_*e*_, *r*, and *L*, rather than the per-site recombination rate alone. Assuming loci of 1,000 nucleotides and an effective size of *N*_*e*_=10, 000, the average human recombination rate, *r*=1.26 cM/Mb (Jensen-Seaman et al., 2004), corresponds approximately to *ρ*=0.5. Using *ρ*=0.5 as a reference, we used two higher values, *ρ*=1.5 and *ρ*=15, corresponding to recombination rates 3 and 30 times higher than the human average. These higher values were chosen to represent organisms with elevated recombination rates (Zhang, 2003; Rizzon et al., 2006), or genomic regions comparable to recombination hotspots (Jeffreys et al., 2001). For each combination of simulation parameters, we also included a non-recombining control with *ρ*=0. The simulation parameters and their fixed or explored values are summarized in Supplementary Table S1. Supplementary Table S2 shows the correspondence between each simulation identifier and its corresponding values of ILS, GF, and recombination settings.

When recombination occurs, `ms` produces one true local tree for each non-recombining segment of the simulated locus. However, `Aphid` takes one tree per locus as input. We therefore summarized the set of local trees generated within each recombining locus into a single locus tree. To do so, each local tree was first transformed into a pairwise distance matrix between species (Fig. 2). We then built a consensus distance matrix in which each pairwise distance corresponded to the mean distance across local trees, weighted by the physical length of the corresponding non-recombining segment. This weighting ensured that longer segments contributed more strongly to the final locus tree than shorter segments. The final consensus distance matrix was then transformed into a single tree using the Neighbor-Joining algorithm implemented in the `bionj()` function of the `phytools` R package (Revell, 2024). This procedure was designed to mimic the practical situation in which a recombining locus is represented by a single inferred gene tree. Treating recombining and non-recombining loci identically isolates recombination from phylogenetic reconstruction error, without adding additional methodological biases.

**Fig. 2.**
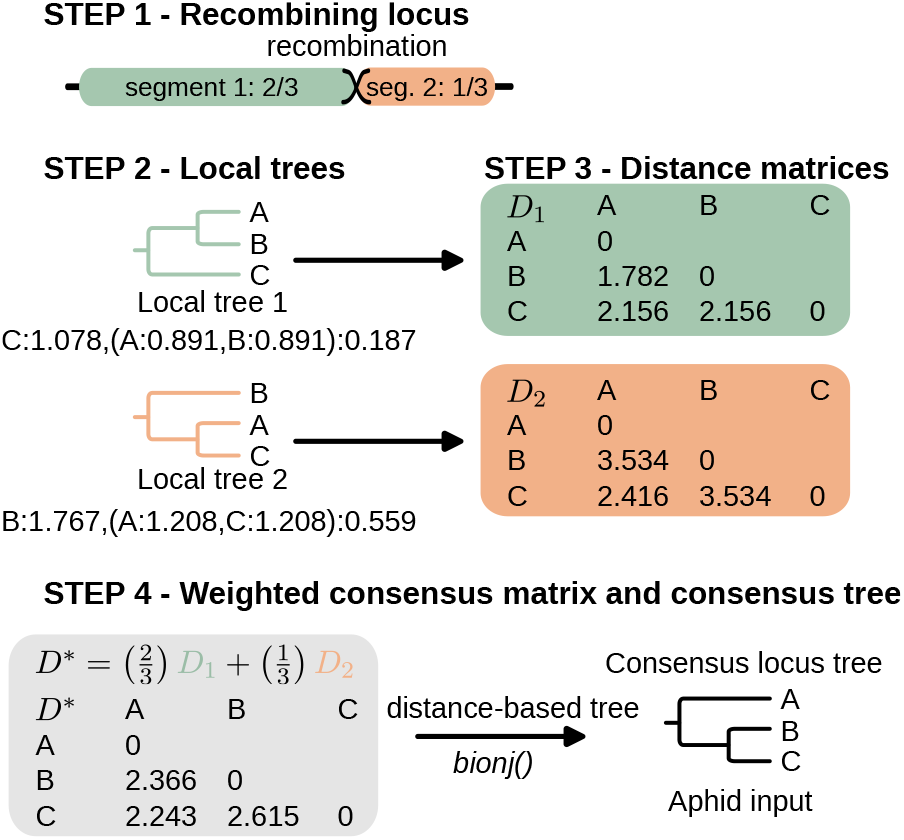
Procedure used to summarize a recombining locus into a single locus tree. When recombination occurs within a locus, `ms` produces one local genealogy for each non-recombining segment. In the example shown here, the locus is composed of two segments representing two-thirds and one-third of the sequence, respectively. Each local tree is first converted into a pairwise distance matrix between species. A consensus distance matrix is then obtained by averaging pairwise distances across local trees, weighted by the physical length of the corresponding segment. This weighted matrix is finally transformed into a single BioNJ tree, which is used as the locus-level input for `Aphid`.

To characterize the genealogical complexity generated by each recombination rate, we recorded the number of non-recombining segments per locus, the topology of their local trees, and the proportions of loci containing multiple local trees or multiple topologies (Supplementary Table S3).

### Aphid Analysis

All datasets were analysed using `Aphid` version 0.11 (Galtier, 2024). For each dataset, the input consisted of one tree per simulated locus, obtained either directly from the non-recombining simulation or from the consensus procedure described above for recombining loci. The taxon file provided to `Aphid` defined *A, B*, and *C* as the focal triplet ((*A, B*), *C*) (Fig. 1), *D, E*, and *F* as additional non-focal species, and *G, H*, and *I* as outgroups. All analyses were performed using the species tree ((*A, B*), *C*) as the reference topology for the focal triplet.

Because branch lengths were rescaled to substitutions per site, many simulated internal branches were shorter than in typical empirical gene trees. By default, `Aphid` considers a triplet as unresolved when the internal branch length, multiplied by the sequence length, is lower than the parameter `min d`, which is set to 0.5. Using this default value caused most simulated trees to be treated as unresolved, not because of genuine topological uncertainty, but because of the small branch lengths imposed by our scaling. We therefore decreased `min d` to 0.0001. This change only affected the threshold used to classify very short internal branches as unresolved; it did not modify the branch lengths or the topology of the input trees.

To evaluate whether `Aphid` outputs contain a usable relative timing signal, we allowed GF coalescence times to vary across ten values, expressed relative to the *A*–*B* divergence time *t*_1_. These values ranged from 0.1 to 1.0, in increments of 0.1, thereby covering GF events from relatively recent to close to the *A*–*B* split. Following the recommendation of Galtier (2024), we excluded replicate datasets in which the concordant topology ((*A, B*), *C*) represented less than 50% of the analysed trees. `Aphid` treats GF and ILS as distinct scenarios, and its approximation becomes difficult to interpret when phylogenetic conflict is so extensive that histories combining GF and ILS are expected to become frequent.

### Performance Assessment

From `Aphid` output, we extracted three quantities for each simulated dataset: the proportion of genes inferred to be affected by ILS or GF, the relative timing of GF, and its direction. Unless otherwise stated, all quantities were computed over the set of loci retained by `Aphid` after filtering, and *n* denotes the number of retained loci.

We first estimated the contribution of GF to phylogenetic conflict. Because empirically the true status for each locus is unknown to `Aphid`, this contribution was calculated as a posterior expectation. For each locus *i, P* (*GF* | *G*_*i*_) can be interpreted as the expected contribution of this locus to the total number of loci affected by GF. Summing these posterior probabilities across loci therefore gives the expected number of loci affected by GF. Let *T* denote one of the two discordant topology classes. For each discordant topology class *T*, the contribution of this class to the dataset-wide inferred contribution of GF was calculated as:

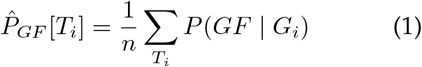

Where *G*_*i*_ is the tree inferred for locus *i, P* (*GF* | *G*_*i*_) is the posterior probability that this locus was affected by GF, and the sum is taken over all loci whose topology belongs to class *T* . For each discordant topology class *T*, this expression corresponds to the topology-specific contribution defined by Galtier (2024). The overall inferred contribution of GF to phylogenetic conflict was obtained by summing 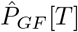 over the two discordant topology classes. The same approach was used to estimate the contribution of ILS:

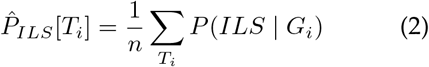

Similarly, the overall inferred contribution of ILS was obtained by summing 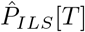 over the two discordant topology classes. To compare the accuracy of 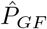 among recombination treatments, we retained simulations with 10% and 25% GF. Simulations with 50% GF were not included because the number of datasets passing the `Aphid` filter differed markedly among values of *ρ*. For each retained dataset, we calculated the bias, defined as 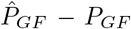, together with the absolute and squared errors. These quantities were analysed using linear models including recombination rate, ILS level, simulated GF proportion, GF timing, and GF direction, as well as the interaction between recombination rate and each of the other factors. Estimated marginal means and contrasts with the non-recombining treatment were obtained with the R package `emmeans` (Lenth and Piaskowski, 2025). RMSE was calculated as the square root of the estimated mean squared error, whereas statistical tests were performed on squared error.

We then estimated the relative timing of GF from the posterior probabilities assigned by `Aphid` to the different GF timing categories. Because GF can generate either of the two discordant topologies, timing was estimated separately for each discordant topology class. Let *T* denote one of these classes, with *T* =*AC* corresponding to the topology ((*A, C*), *B*), and *T* =*BC* corresponding to the topology ((*B, C*), *A*). For each discordant topology class *T*, we considered the set *D*_*T*_ of retained loci whose inferred topology belonged to *T*, and denoted its size by *n*_*T*_ . For each locus *i* ∈ *D*_*T*_, we first calculated a conditional timing estimate as the posterior-weighted mean of the GF timing categories:

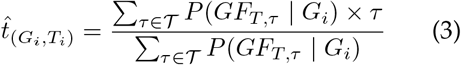

Where *T* = *{*0.1, 0.2, …, 1.0*}* is the set of GF times tested in `Aphid`, expressed relative to *t*_1_ (Fig. 1), and 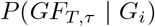 is the posterior probability that locus *i* supports a GF scenario associated with topology *T* and time *τ* .The topology-specific timing estimate was then calculated as a GF-posterior-weighted mean across discordant loci:

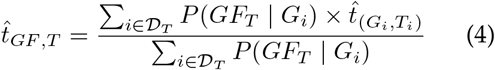

This definition separates the amount of GF inferred in a dataset from the inferred timing of GF: loci with low posterior support for GF contribute little to the timing estimate. To quantify timing accuracy, we retained simulations with 10% and 25% GF and used 0.1, 0.5, and 0.9 (relative to *t*_1_) as the reference timings for recent, intermediate, and ancestral GF, respectively. A timing estimate was considered estimable when it was defined and strictly greater than zero. For estimable replicate datasets, we calculated the bias, defined as the inferred minus the reference timing, together with the absolute and squared errors. These quantities were analysed using the same linear-model framework as for the inferred proportion of GF. Because failures to estimate timing represent a distinct component of performance, the proportion of replicate datasets producing an estimable timing was analysed separately.

Finally, we evaluated whether `Aphid` recovered the simulated direction of GF. For each replicate dataset, we calculated the mean posterior support for the two possible directions of GF between *A* and *C*, using the *n*_*d*_ retained loci whose inferred tree was discordant with the species tree.

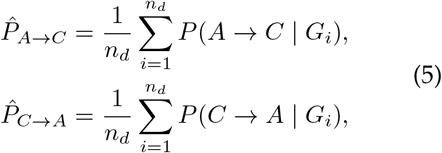

where *P* (*A* → *C* | *G*_*i*_) and *P* (*C* → *A* | *G*_*i*_) are the posterior probabilities that the discordant genealogy of locus *i* was generated by GF in the corresponding direction.

Directional support was then summarized using the Bayes factor:

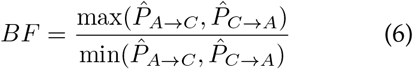

For each retained replicate dataset with GF, the direction with the highest posterior support was recorded as the preferred direction. Replicate datasets without a defined preferred direction were excluded when calculating the frequency with which the preferred direction matched the simulated direction.

The strength of directional support was assessed separately. A preferred direction was considered supported when *BF* ⩾10. Supported directions were classified as correct when they matched the simulated direction and as wrong otherwise. Replicate datasets with *BF*<10, or for which the Bayes factor was undefined, were classified as ambiguous. Both the frequency of a correct preferred direction and the frequencies of ambiguous, supported-correct, and supported-wrong outcomes were summarized separately by recombination rate and simulated GF direction.

### Inference of Gene Flow Timing in African Cichlids

We reanalysed the exon dataset of Astudillo-Clavijo et al. (2023) to characterize the relative timing of the previously inferred GF signal involving *Coptodon* and closely related West and Central African cichlid lineages. We downloaded the 588 exon alignments used in the original study, together with the corresponding locus information from the original Table S2. Because the alignments could contain non-coding sequence and did not necessarily start in the correct reading frame, we used the *Oreochromis niloticus* GenBank accession provided for each exon to retrieve the corresponding annotated transcript and CDS coordinates. These coordinates were projected onto the exon alignments to identify the coding region and codon positions. After filtering loci for which the CDS could not be reliably identified or the reading frame was ambiguous, 517 exon alignments were retained. For each locus, we first inferred the gene-tree topology from the complete CDS, using all three codon positions, with IQ-TREE (Wong et al., 2026) and model selection with ModelFinder (Kalyaanamoorthy et al., 2017). Once the topology was fixed, branch lengths were re-estimated using third codon positions only, thereby reducing the contribution of non-synonymous substitutions to the branch-length signal used by `Aphid`. The locus length supplied to `Aphid` was consequently defined as the number of third-codon-position sites used to estimate these branch lengths. Gene trees were rooted using the non-cichlid outgroups available for each locus; 61 loci without a usable outgroup were excluded, leaving 456 rooted gene trees. The rooted 588-gene ASTRAL species tree from Astudillo-Clavijo et al. (2023) was used as the reference species tree for defining the focal triplets. We focused on the GF signal involving *Coptodon* identified in the original network analysis and repeated the analysis with four representatives of this genus (*C. zillii, C. guineensis, C. bakossiorum*, and *C. flavus*). For each *Coptodon* species, focal triplets included *Gobiocichla ethelwynnae* and one of five taxa from the putatively affected lineage: *Congolapia bilineata, C. crassa, Chilochromis duponti, Steatocranus gibbiceps*, or *S. rouxi*. Crossing these five taxa with the four *Coptodon* species resulted in 20 focal triplets (Supplementary Table S11). Twelve additional triplets involving *Gobiocichla, Pelmatolapia mariae*, and *Heterotilapia buttikoferi* were used as controls, resulting in 32 triplets in total. Depending on taxon coverage, between 209 and 228 of the 456 gene trees were available for each triplet. For each comparison, we estimated the proportion of loci attributed to GF 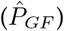 and its relative timing 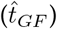 as described above. When the same species pair occurred in several triplets, 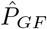 was averaged across triplets for visualization.

## Results

### Effect of Recombination on Gene-Flow Proportion Estimates

We first quantified the effect of intra-locus recombination on the proportion of loci inferred by `Aphid` as affected by GF, across ILS levels, simulated proportions of migrant loci, and GF timings. For each combination of parameters, we calculated the mean inferred proportion across retained replicate datasets (Fig. 3).

**Fig. 3.**
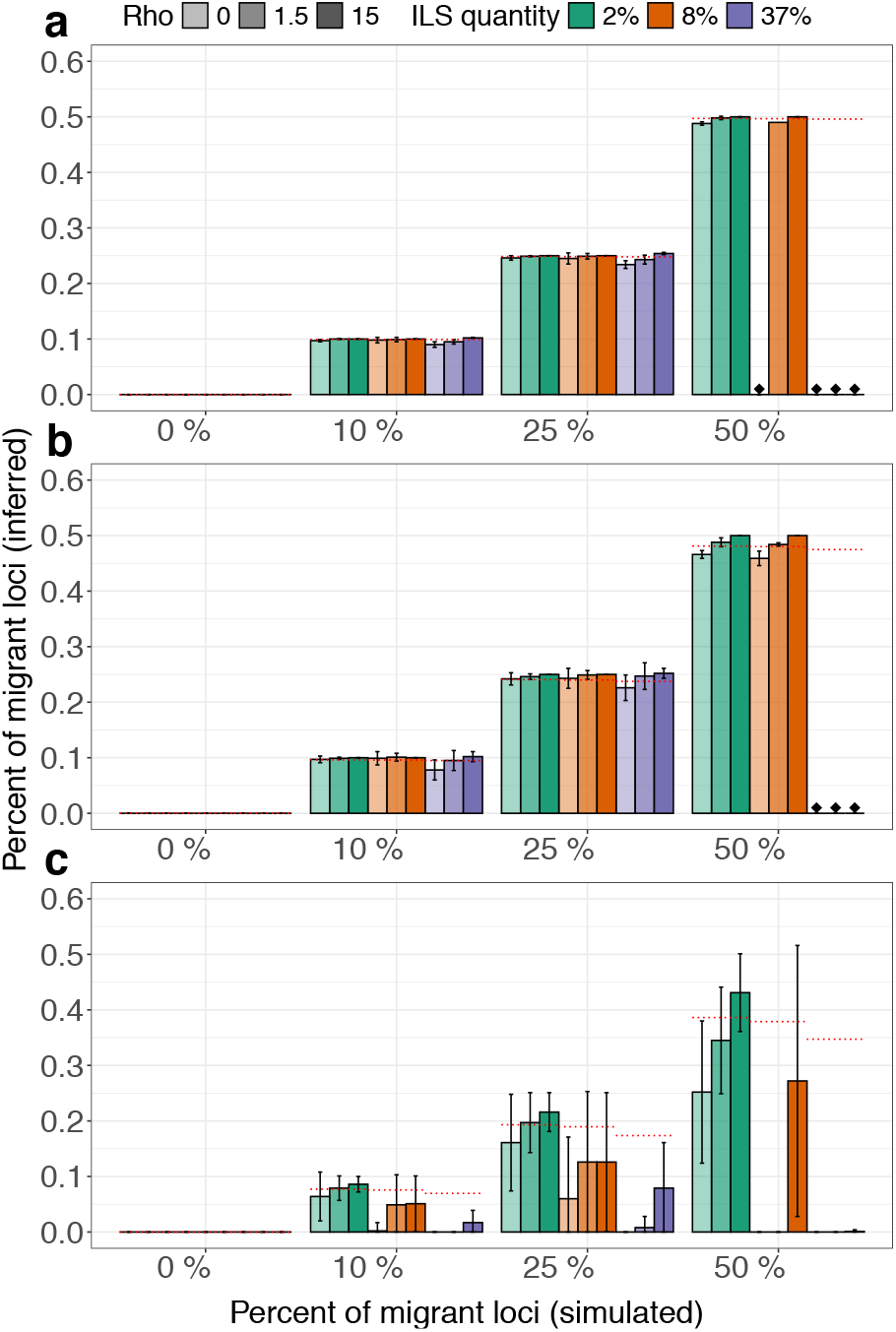
Inferred proportion of loci affected by gene flow 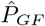, equation 1) between non-sister species, for recent GF (a), intermediate GF (b) and ancestral GF (c). Bars represent the mean proportion of loci inferred by `Aphid` as affected by GF. Error bars show ± one standard deviation across retained replicate datasets. Colours indicate thone standard deviation across retained replicate datasets. Colours indicate the level of ILS: green for 2%, orange for 8%, and blue for 37%. Colour intensity indicates the recombination rate, from the lightest shade for *ρ* = 0, to the intermediate shade for *ρ* = 1.5, and the darkest shade for *ρ*=15. Red dashed lines indicate the expected values under each simulated scenario, accounting for the proportion of loci with discordant topology generated by ILS. Black diamonds indicate parameter combinations for which no replicate dataset satisfied the requirement that at least 50% of analysed trees have the concordant topology. Means were calculated across retained replicate datasets; the number of replicates retained for each affected parameter combination is reported in Supplementary Table S4.

When no GF was simulated, 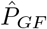 remained at or very close to zero across ILS and recombination treatments (Fig. 3). Thus, increasing intralocus recombination did not increase the inferred GF contribution in simulations without GF.

When recent GF was simulated, `Aphid` recovered proportions close to the expected values across most parameter combinations (Fig. 3a). Among retained simulation conditions, the maximum underestimation was 0.014. The 50% concordance criterion mainly excluded replicate datasets with 50% GF, particularly under high ILS, because these datasets frequently contained a majority of discordant trees. The proportion of replicate datasets satisfying the requirement that concordant trees represent at least 50% of the analysed loci is reported in Supplementary Table S4 (combinations not listed have passed this criterion in all 100 replicates).

Intermediate GF showed the same general pattern, although variation among replicate datasets was slightly greater than for recent GF (Fig. 3b). Across retained simulations, deviations from the expected proportions remained limited, ranging approximately from − 0.02 to 0.02. The 50% concordance criterion again mainly excluded replicate datasets with 50% GF under high ILS. Ancestral GF behaved differently (Fig. 3c).

With 2% ILS, inferred proportions remained close to the expected values, although they were slightly overestimated in some recombining simulations. As ILS increased, however, the inferred proportion of GF decreased and variation among replicate datasets increased. This effect was strongest with 37% ILS, where the inferred proportion was often close to zero, indicating that ancestral GF became difficult to distinguish from ILS.

Across retained simulations with 10% and 25% GF, and averaged over ILS level, GF timing, proportion, and direction, the mean bias was − 0.046 at *ρ*=0, − 0.032 at *ρ*=1.5, and − 0.025 at *ρ*=15. This reduction in bias and estimation error with increasing *ρ* was strongest for ancestral GF, whereas estimates for recent and intermediate GF remained close to the simulated proportions (Supplementary Tables S5 and S6).

These patterns were observed despite a strong increase in within-locus genealogical complexity. Loci contained on average ∼ 90 non-recombining segments at *ρ*=1.5, compared with *∼* 600 at *ρ*=15, and the proportion containing multiple topologies increased from 0.20–0.23 to 0.51–0.63 (Supplementary Table S3). Thus, increasing recombination did not reduce the accuracy of the inferred GF proportion under the conditions tested. The main limitation remained the inference of ancestral GF under high ILS, where GF was strongly underestimated. These analyses combine both directions of GF; direction-specific results are provided in Supplementary Figs. S1 and S2.

### Effect of Recombination on the Inference of Gene Flow Timing

We then asked whether the branch-length signal retained by `Aphid` contained information about the relative timing of GF. For each replicate dataset, we summarized the posterior support across loci and recorded the GF timing category receiving the highest support. For each combination of parameters, we then calculated the pro-portion of replicate datasets assigned to each timing category. Results for simulations in which 10% of loci were affected by GF from species *C* to *A* are shown in Fig. 4, and results for GF from *A* to *C* are shown in Fig. 5.

**Fig. 4.**
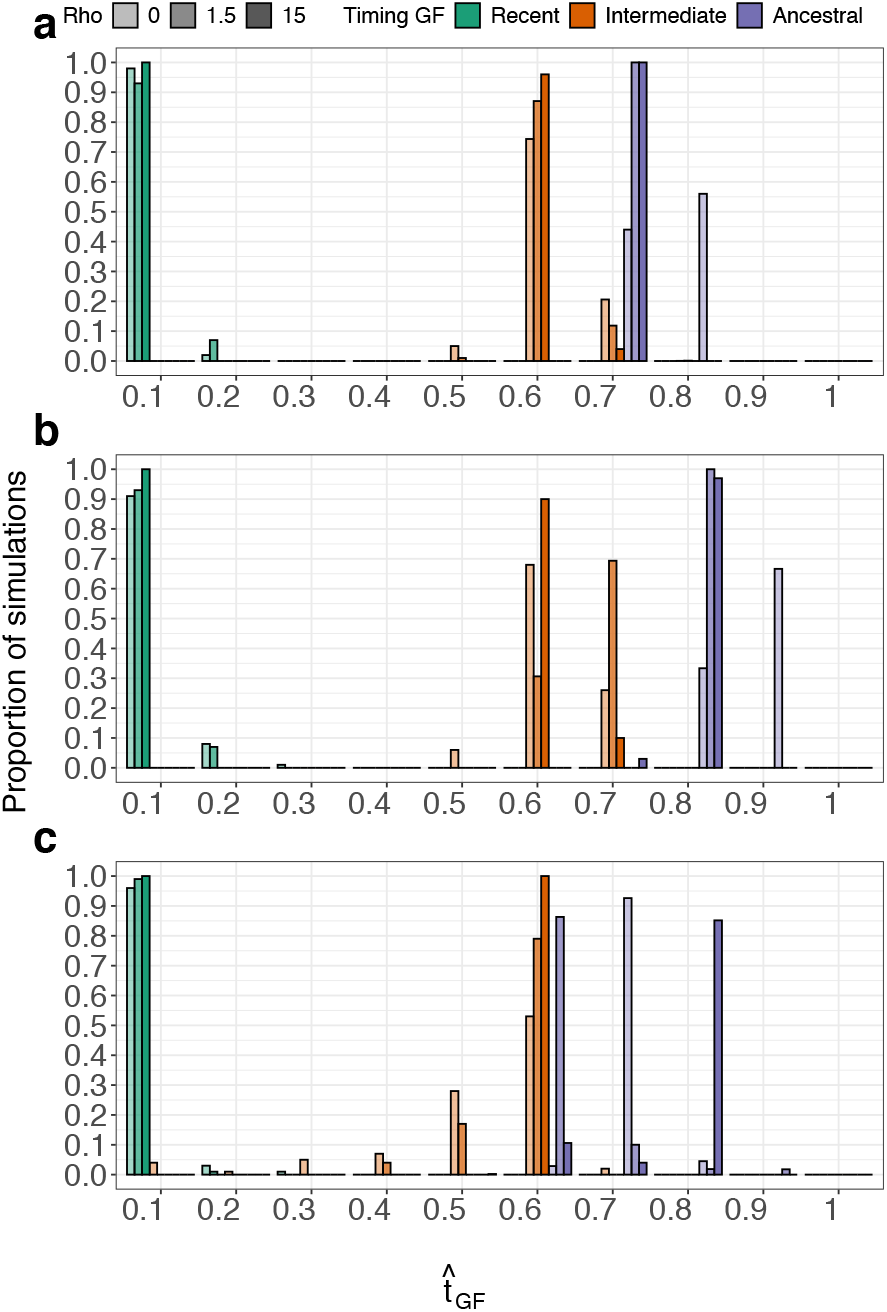
Timing inferred by `Aphid` 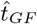, equation 4) for simulation in which 10% of loci were affected by GF from species *C* to species *A*. Panels correspond to the three levels of ILS: 2% in panel (a), 8% in panel (b), and 37% in panel (c). Bars represent the proportion of replicate datasets assigned to each inferred GF timing category. Colours indicate the simulated timing of GF: green for recent GF, orange for intermediate GF, and blue for ancestral GF. Colour intensity indicates the recombination rate, from the lightest shade for *ρ*=0, to the intermediate shade for *ρ*=1.5, and the darkest shade for *ρ*=15.

**Fig. 5.**
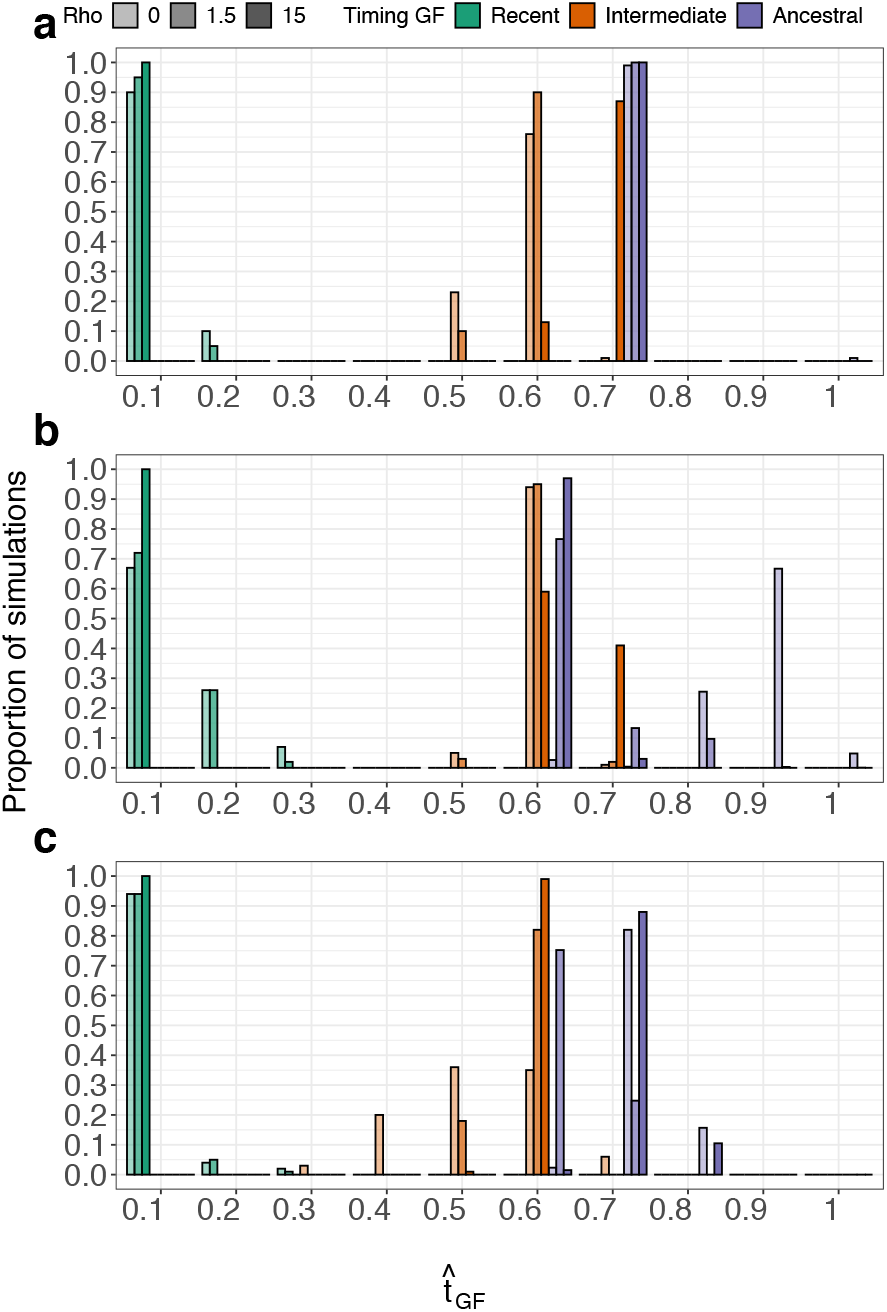
Timing of GF inferred by `Aphid` for simulation in which 10% of loci were affected by GF from species A to species C. Panels, bars, colours, and colour intensities are as in Fig. 4

Recent GF was consistently inferred as recent. For *C* → *A* GF, this pattern was observed across recombination rates and ILS levels (Fig. 4, green bars), and the same result was obtained for *A* → *C* GF (Fig. 5). Quantitatively, timing error was already small without recombination and did not increase as *ρ* increased. The MAE was 0.0095 at *ρ*=0, 0.0074 at *ρ*=1.5, and 0 at *ρ*=15 (Supplementary Table S7).

Intermediate GF was generally inferred as older than simulated. For *C* → *A* GF, most replicate datasets were assigned to timing categories between 0.6 and 0.8 (Fig. 4, orange bars). The shift towards older values increased with recombination: the mean bias was 0.075 at *ρ*=0, 0.101 at *ρ*=1.5, and 0.121 at *ρ*=15. Correspondingly, the MAE increased from 0.086 without recombination to 0.102 and 0.121 in the two recombining treatments (Supplementary Table S7). This shift increased the overlap between intermediate and ancestral timing distributions in some parameter combinations.

Ancestral GF showed the opposite bias and was generally inferred as more recent than simulated. For *C* → *A* GF, most inferred timings fell between 0.7 and 1.0, except under high ILS, where they frequently shifted towards intermediate values between 0.3 and 0.7 (Fig. 4, blue bars). The mean bias was − 0.144 at *ρ*=0, − 0.201 at *ρ*=1.5, and − 0.170 at *ρ*=15. Recombination therefore increased the magnitude of the error, although not monotonically: the MAE increased from 0.147 at *ρ*=0 to 0.210 at *ρ*=1.5 and 0.187 at *ρ*=15 (Supplementary Table S7).

The *A* → *C* simulations gave a broadly similar picture (Fig. 5). Recent GF was again inferred accurately, whereas intermediate GF tended to be inferred as older and ancestral GF as more recent than simulated. Intermediate and ancestral timings were generally more clearly separated than for *C* → *A* GF, although ancestral GF again shifted towards intermediate values under high ILS (Fig. 5c). The magnitude of the recombination effect varied significantly with both ILS level and GF direction, as well as with the simulated timing and proportion of GF (Supplementary Table S8).

Timing was estimable in all replicate datasets with recent or intermediate GF. Failures were restricted to ancestral GF and differed among recombination treatments. An ancestral timing was estimable in 89.6% of replicate datasets at *ρ*=0, 99.8% at *ρ*=1.5, and 92.6% at *ρ*=15 (Supplementary Table S9). Accuracy estimates for ancestral GF are therefore conditional on the replicate datasets in which a positive timing estimate was obtained.

Finally, these analyses show that `Aphid` retains a strong relative timing signal for recent GF. Intermediate and ancestral GF remain partly distinguishable, but both exhibit systematic errors in opposite directions: intermediate GF is inferred as relatively too old, whereas ancestral GF is inferred as relatively too recent. Intra-locus recombination does not reduce timing accuracy for recent GF, but increases error for intermediate and ancestral GF, with an effect that depends strongly on ILS level and GF direction. Results for simulations in which 25% and 50% of loci were affected by GF showed similar distributions and are provided in Supplementary Figs. S3–S6.

### Effect of Recombination on the Inferred Direction of Gene Flow

We then examined whether intra-locus recombination affected the directional information contained in `Aphid` outputs. Directional support was generally weak. More than 98% of retained replicate datasets were classified as ambiguous at every recombination rate and for both simulated directions because the Bayes factor was lower than 10 or undefined (Supplementary Table S10). Strong support for the correct direction never exceeded 0.4% of replicate datasets within a recombination treatment and simulated direction. Supported but incorrect directions were restricted to simulations of *C* → *A* GF at *ρ*=0, where they represented 1.12% of replicate datasets.

Despite this weak absolute support, among datasets for which a preferred direction was defined, the simulated direction received the higher posterior support more often than the alternative direction at every recombination rate (Table 1).

**Table 1.** Percentage of retained replicate datasets in which the simulated direction of GF received the highest posterior support. Denominators are the numbers of retained replicate datasets with a defined preferred direction: *n*=1686, 1917, and 2398 for simulated *A→C* at *ρ*=0, 1.5, and 15, respectively, and *n*=2039, 2172, and 2234 for simulated *C→A*.

| $\rho$ | Simulated $A \rightarrow C$ | Simulated $C \rightarrow A$ |
| --- | --- | --- |
| 0 | 75.33% | 81.31% |
| 1.5 | 79.13% | 82.64% |
| 15 | 83.57% | 86.84% |

The frequency with which the correct direction ranked first increased with *ρ* for both simulated directions (Table 1).

Increasing recombination therefore made the correct direction more likely to receive the highest posterior support, but did not produce strong evidence allowing that direction to be inferred confidently.

### Relative Timing of Gene Flow in African Cichlids

We used gene-tree branch lengths to assess whether the *Coptodon*-associated reticulation identified by Astudillo-Clavijo et al. (2023) was compatible with recent or older GF. Across the 20 focal cichlid triplets, 28.3–30.6% of the retained gene trees were discordant with the species-tree topology, with discordant trees supporting several alternative relationships (Fig. 6a). For the discordant topology grouping the focal taxon with *Coptodon*, `Aphid` inferred 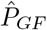 values ranging from 0.017 to 0.045 (Fig. 6b). Within this focal discordant topology, 9.7–25.1% of the signal was attributed to GF rather than ILS (Supplementary Table S12). GF contributions were generally higher than in the control comparisons, where 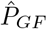 ranged from 0.004 to 0.026. The signal was particularly consistent for *Congolapia* and *Chilochromis*, for which estimates ranged from 0.028 to 0.045 across the four *Coptodon* species.

**Fig. 6.**
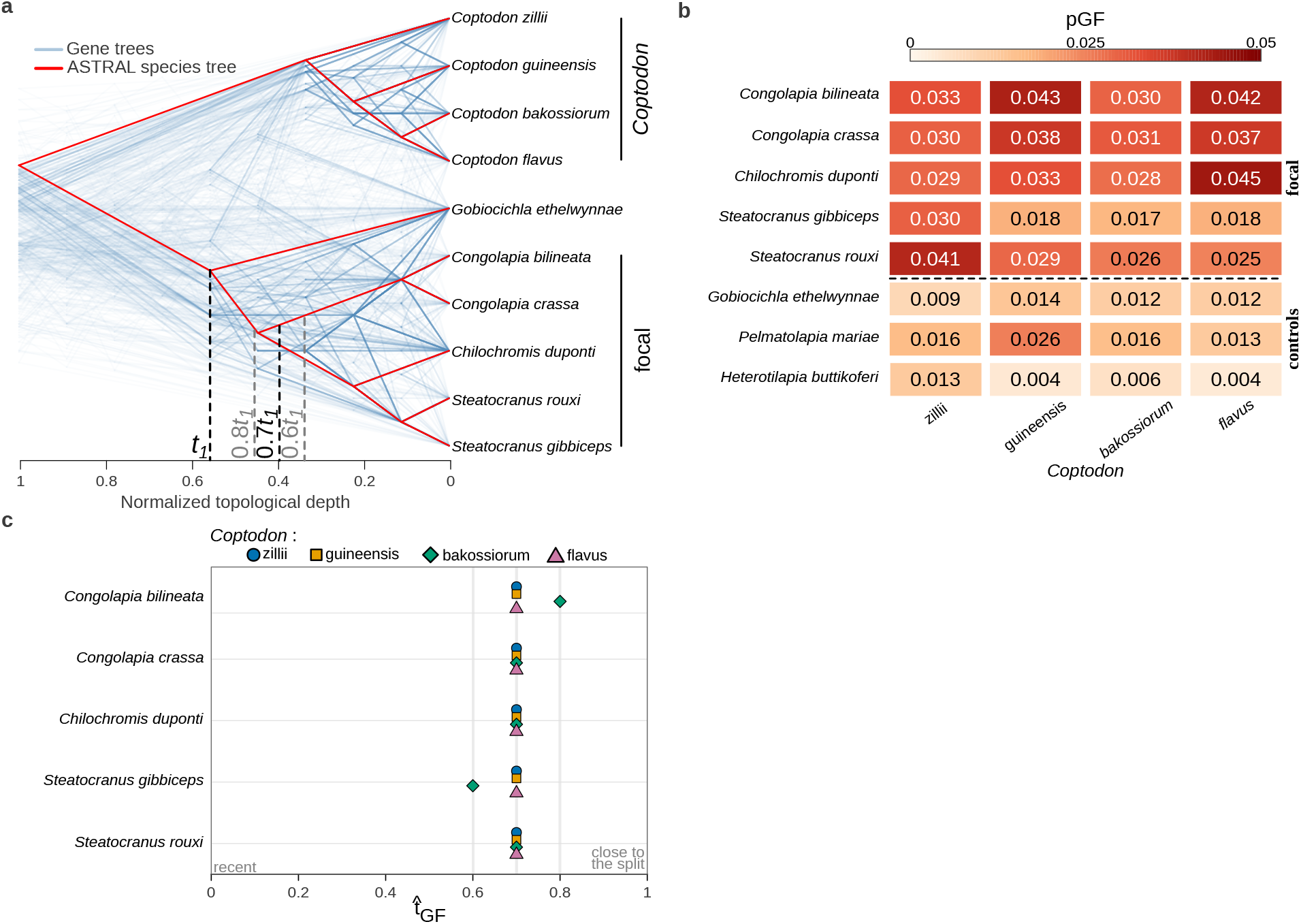
Empirical application of `Aphid` to an African cichlid dataset comprising 456 exon gene trees. (a) DensiTree representation of gene trees discordant with the rooted ASTRAL species tree for the ten taxa included in the focal phylogenetic comparison. Gene trees are shown in blue and the species tree in red. Trees were rescaled to a normalized topological depth for visualization; horizontal positions therefore do not represent molecular branch lengths or divergence times. Dashed vertical lines indicate *t*_1_ and the 0.6*t*_1_, 0.7*t*_1_, and 0.8*t*_1_ inferred relative timings shown in panel (c). (b) Estimated proportion of loci affected by gene flow 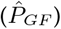 between each row taxon and four *Coptodon* species. The upper five rows correspond to the focal comparisons, whereas the lower three rows are controls. (c) Relative timing of GF-associated coalescence 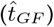 for the focal comparisons. Values are expressed relative to *t*_1_, with 0 indicating recent coalescence and 1 indicating coalescence close to the split between the sister lineages of the corresponding triplet. Symbols and colours identify the four *Coptodon* species.

Despite variation in the inferred amount of GF, its relative timing was remarkably consistent across focal comparisons. Eighteen of the 20 comparisons yielded 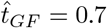, with the two remaining estimates at 0.6 and 0.8 (Fig. 6c). Within each focal taxon, replacing *C. zillii* with the three alternative *Coptodon* representatives changed 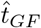 by at most 0.1. Thus, all inferred GF-associated coalescence times were concentrated between 0.6 and 0.8 of the interval from the present to the sister-lineage split, with none supporting a recent timing.

Our simulations show that estimates in this range cannot reliably distinguish intermediate from ancestral GF, because intermediate events tend to be inferred as too old whereas ancestral events tend to be inferred as too recent. The cichlid analysis therefore does not provide a precise timing for the proposed GF involving *Coptodon*, but consistently rejects the hypothesis of recent GF.

## Discussion

Our simulations show that the single-tree approximation is less fragile than might be expected, although its robustness differs among components of the inference. Summarizing several local genealogies into one locus tree did not reduce the accuracy of the inferred proportion of loci affected by GF. Instead, recombination reduced the average underestimation of this proportion, particularly for ancestral GF. However, the relative timing signal was more sensitive. Recent GF remained accurately identified, whereas recombination increased timing error for intermediate and ancestral events. Directional inference remained accurate but weak, suggesting that the information required to estimate the prevalence, timing, and direction of GF is progressively more fragile.

The absence of a detrimental effect of recombination on the inferred proportion of GF is consistent with a simple coalescent expectation. Recombination changes how genealogies are correlated along a locus, but it does not change the marginal distribution of a local genealogy taken at a single position. If `Aphid` could be applied directly to true local trees, recombination alone would therefore not be expected to create a systematic bias. The practical issue is that empirical analyses do not use all local trees separately: they usually reduce each locus to one inferred tree. Our simulations show that this reduction did not, by itself, strongly distort the branch-length signal used by `Aphid` . Consistent with this result, 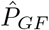 remained close to zero in simulations without GF across recombination treatments, indicating that recombination alone did not generate an appreciable inferred GF contribution. A likely explanation is that, for the locus length and recombination rates explored here, the consensus tree often retained the dominant genealogical signal of the locus, so that recombination changed the local history of some segments without fully erasing the information carried by the locus as a whole. This interpretation is consistent with simulations reported for `QUIBL`, another branch-length-based method, for which inference remained robust to recombination until recombination substantially exceeded the mutation rate (Edelman et al., 2019). These simulations suggest that reducing a recombining locus to a single tree need not bias the inferred prevalence of GF, provided that the locus-level tree retains its dominant genealogical signal. This conclusion, however, does not extend unchanged to the inferred timing of GF.

The amount of genealogical heterogeneity within a locus depends not only on the per-site recombination rate, but also on the scale at which loci are defined. Because *ρ*=4*N*_*e*_*rL*, increasing locus length *L*, recombination rate *r*, or effective population size *N*_*e*_ increases the expected amount of recombination within a locus. Our simulations kept locus length fixed at 1,000 nt and varied *ρ* through *r*, so they isolate the consequences of increasing within-locus recombination but do not capture the additional phylogenetic information provided by longer sequences. This creates a practical trade-off in locus definition: shorter loci reduce the opportunity for intra-locus recombination, whereas longer loci contain more informative sites and can reduce gene-tree estimation error. Our results therefore do not identify an optimal locus length, but emphasize that recombination should be considered at the scale of the loci used for phylogenetic inference rather than from the per-site recombination rate alone.

Ancestral GF remained the most difficult scenario to infer correctly in our analyses. As GF is moved deeper in the history of the triplet, the branch-length contrast separating GF from ILS progressively erodes. Lineages affected by ancestral GF and those affected by ILS both trace back to deeper ancestral populations, causing their branch-length distributions to overlap. High ILS accentuates this problem by increasing the proportion of discordant genealogies with deep coalescence times. This explains why the proportion of loci affected by ancestral GF was underestimated, particularly under high ILS, regardless of recombination treatment. This loss of temporal resolution is not specific to `Aphid` . Populationgenomic demographic approaches such as `DILS` and ∂*a*∂*i* can also model ancestral migration, but the timing of older migration events can likewise be weakly identifiable and confounded with other demographic parameters (Fraïsse et al., 2021; Gutenkunst et al., 2009; Sousa et al., 2011).

Recombination nevertheless contributed an additional and distinct source of error in the timing inference. Intermediate GF was systematically inferred as older than simulated, whereas ancestral GF was inferred as more recent. These opposite biases effectively compress the temporal signal, causing intermediate and ancestral GF to converge towards overlapping inferred time ranges. Both shifts increased with recombination, although the effect on ancestral GF was not monotonic with *ρ* and depended strongly on ILS level and GF direction. Thus, the poor recovery of ancestral GF cannot be attributed exclusively to recombination, but neither can recombination be considered neutral for its inferred timing. The intrinsic overlap between ancestral GF and ILS limits the available branch-length information, while the reduction of multiple local genealogies to a single locus tree further alters how this remaining timing signal is represented.

Despite these biases, the branch-length signal retained substantial information about the relative timing of GF. Recent events were consistently recovered as recent and showed no loss of accuracy with increasing recombination. Intermediate and ancestral events also remained partly distinguishable, but their inferred timings converged: intermediate GF shifted towards older values, whereas ancestral GF shifted towards more recent values. Consequently, timing estimates should not be interpreted as unbiased estimates of an absolute date. They are more appropriately viewed as a relative signal that reliably separates recent from older GF, but provides less resolution within the older part of the species history.

Our cichlid results refine rather than challenge the interpretation of Astudillo-Clavijo et al. (2023). Their SNaQ analysis of former-tilapiines inferred a reticulation involving *Coptodon zillii* and the Tilapiini–Steatocranini lineage, with an inheritance probability of approximately 0.30. More generally, they concluded that hybridization contributed to specific phylogenetic conflicts, whereas ILS remained a prevailing source of gene-tree incongruence across African cichlids. Our branch-length analysis is consistent with this interpretation: despite substantial gene-tree discordance, only 9.7–25.1% of the signal supporting the focal discordant topology was attributed to GF rather than ILS. These estimates should not be compared directly with the SNaQ inheritance probability, because the two quantities have different definitions.

What our analysis adds is information about the relative timing of the previously identified GF signal. Across the 20 focal comparisons, GF-associated coalescence was consistently inferred at 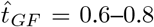, with very little variation when *C. zillii* was replaced by other *Coptodon* representatives. The signal was therefore not restricted to the particular species used to represent these lineages in the original network analysis. Our simulations prevent us from distinguishing intermediate from ancestral GF within this range, but they show that such values are inconsistent with a strongly recent signal. The *Coptodon*-associated reticulation identified by AstudilloClavijo et al. (2023) can therefore be given an additional temporal interpretation: its branch-length signal is more consistent with older than with recent exchange. It remains unclear whether this pattern results from a single ancestral event or from several older episodes of GF.

The distinction between recent and older GF is also relevant to comparative studies of how rapidly species barriers become impermeable. Population genomic approaches can address this question more directly when resequencing data are available, as illustrated by recent comparative analyses based on `DILS` (Fraïsse et al., 2021; Monnet et al., 2025). Phylogenomic datasets, however, are available across a much broader range of taxa. The timing information extracted by `Aphid` could therefore extend comparative analyses of introgression dynamics to clades lacking population-level sampling, provided that analyses emphasize broad temporal contrasts rather than precise estimates of older GF timing.

Directional information showed a different response to recombination. Increasing *ρ* made the correct direction more likely to receive the highest posterior support, indicating that summarizing recombining loci into single trees did not systematically reverse the directional signal. This relative preference was nevertheless too weak to support confident inference. The Bayes factor remained below 10 in almost all replicate datasets, and increasing recombination did not consistently increase the frequency of strongly supported correct directions. Recombination therefore affected the ranking and strength of directional models differently: the correct model ranked first more often, but its advantage over the alternative model remained small.

This weak support is not unexpected. Detecting GF mainly requires showing that a discordant genealogy is shorter than expected under ILS. Inferring direction is harder because it requires the branch-length asymmetry between the two non-sister lineages to be sufficiently strong and consistent to distinguish the two directional scenarios. Our results suggest that this asymmetry was partly retained after recombining genealogies were summarized into a single locus tree, but rarely strongly enough to provide decisive evidence. This is consistent with previous work showing that triplet- or quartet-based approaches often have limited power to infer the direction of introgression from branch-length information alone (Hibbins and Hahn, 2019; Forsythe et al., 2020). Larger phylogenetic contexts may provide additional information, for instance through the relative placement of gene trees within a broader species tree (Mishra et al., 2026). Accordingly, the direction favoured by `Aphid` may provide a weak indication, but should not be interpreted as a supported directional inference without independent evidence.

Our simulation design was deliberately conservative. By working directly from simulated genealogies, we could isolate the effect of intralocus recombination without adding sequence evolution, finite-site effects, alignment error, or phylogenetic reconstruction uncertainty. This was useful for asking whether recombination alone can bias `Aphid`, but it also means that our simulations do not capture the full complexity of empirical datasets. In real phylogenomic analyses, the tree representing a locus may be affected not only by recombination, but also by substitution-rate heterogeneity, model misspecification, limited phylogenetic information, and reconstruction error. These sources of uncertainty may also affect the cichlid analysis, reinforcing the need to interpret its timing estimates as a broad distinction between recent and older GF rather than as precise estimates of when gene exchange occurred. We also considered a single species-tree configuration, constant population size, and one pulse of unidirectional GF between two non-sister lineages. More complex histories, including repeated introgression, bidirectional exchange, variation in population size, or heterogeneous substitution rates, may further blur the branch-length signal used by `Aphid` . Our conclusions should therefore be read as applying to a controlled setting in which recombination is the only source of disagreement between local genealogies within a locus. Timing accuracy was also evaluated conditionally on a positive timing estimate. Although timing was always estimable for recent and intermediate GF, failures occurred for ancestral GF and were unevenly distributed among recombination treatments. The reported error measures therefore describe accuracy when an estimate was available and should be considered together with the probability of obtaining such an estimate. They do not imply that empirical datasets can ignore recombination, but they suggest that intra-locus recombination alone is unlikely to be the dominant source of bias when loci are short enough to retain a clear genealogical signal. Under the conditions explored here, the clearest cost of intra-locus recombination was therefore a loss of temporal resolution for older GF, rather than the creation of spurious GF signal or a systematic distortion of its inferred prevalence.

## Data Availability Statement

The scripts used to perform the simulations, process the trees, run `Aphid`, and reproduce the analyses are available at https://github.com/Arthur-Boddaert/impact-recombination-phylogenetic-inference. Used version of Aphid is available at https://gitlab.mbb.cnrs.fr/ibonnici/aphid/-/tree/e53e0bc88e7f2c2bc648b7f3feb88f1edff4629c/.

## Acknowledgments

We thank Guillaume Achaz, Fabien Condamine and Xavier Vekemans for suggesting this study, and Benoit Nabholz, Christelle Fraïsse, and Nicolas Galtier for their discussions and advice throughout the project.

## Supplementary Materials

### Simulation Design

We summarize here the full simulation design used to evaluate the effect of intra-locus recombination on `Aphid` inference. Table S1 defines the fixed parameters and the values explored, whereas Table S2 provides the complete correspondance between simulation identifiers and parameter combinations.

**Table S1.**
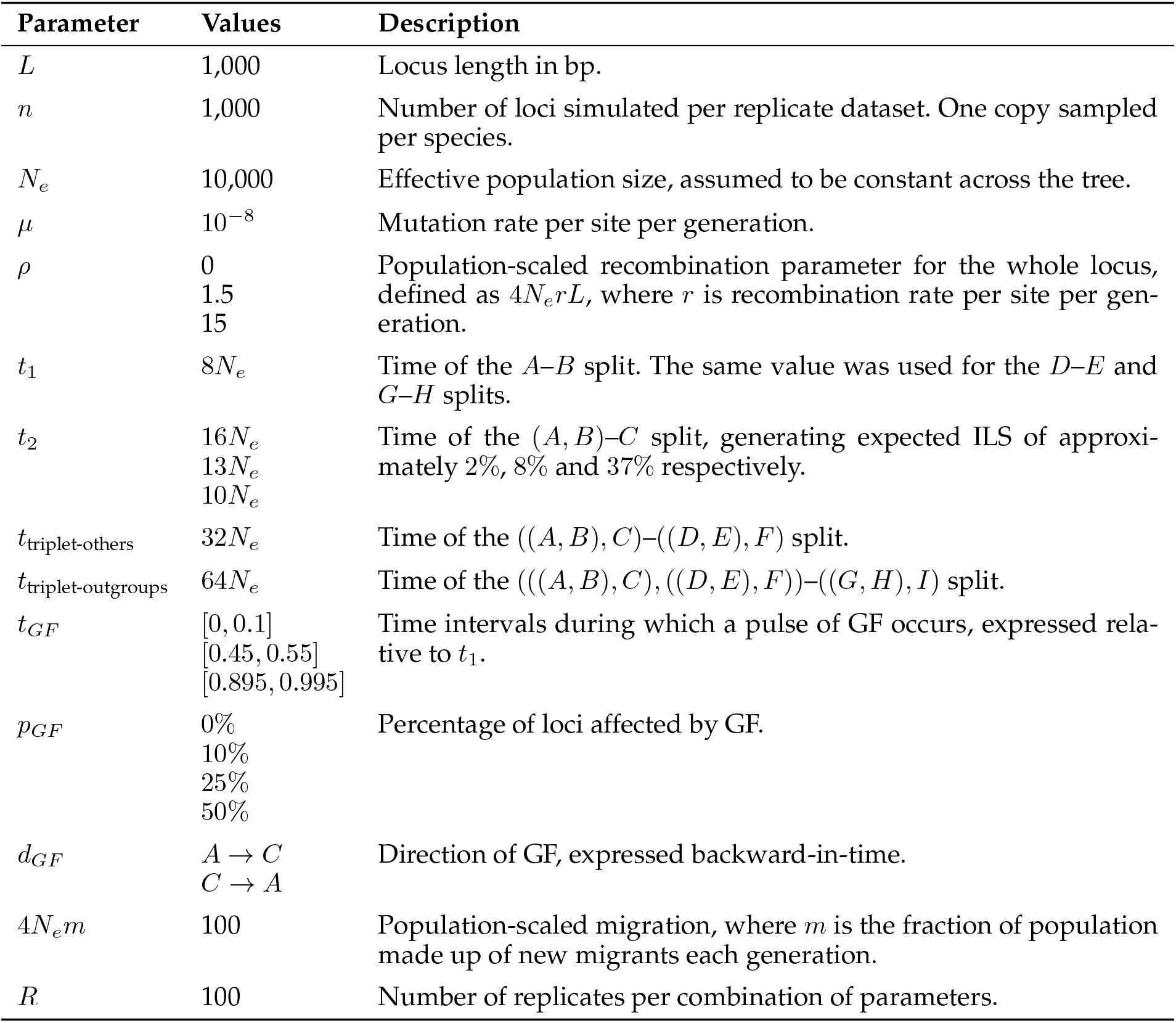
Parameters used to generate the simulated datasets. Fixed values and levels varied among simulation sets are reported together with their definitions. Divergence times are expressed in generations; GF timing windows are expressed relative to *t*_1_, and GF direction is defined backward in time.

### Implementation of Coalescent Simulations

Coalescent simulations were performed with `ms` . Species were numbered from 1 to 9 in the `ms` command lines, corresponding to species *A* to *I* in the main text. Thus, species 1, 2 and 3 correspond to the focal triplet *A, B* and *C*; species 4, 5, and 6 to the additional non-focal species *D, E* and *F* ; and species 7, 8 and 9 to the outgroups *G, H* and *I*.

The following command was used for simulations in which gene flow was simulated from species *C* to *A* backward-in-time with `ms` :

~~~
ms 9 1000 -r <rho> 1000 -T -I 9 1 1 1 1 1 1 1 1 1 0 -em <GF_start> 1
 3 0 -em <GF_start> 3 1 100 -em <GF_end> 1 3 0 -em <GF_end> 3 1 0
 -ej 2 2 1 -ej <t_b> 3 1 -ej 2 5 4 -ej 4 6 4 -ej 2 8 7 -ej 4 9 7
 -ej 8 4 1 -ej 16 7 1
~~~

The following command was used for simulations in which GF was simulated from species *A* to *C* backward-in-time with `ms` :

~~~
ms 9 1000 -r <rho> 1000 -T -I 9 1 1 1 1 1 1 1 1 1 0 -em <GF_start> 3
 1 0 -em <GF_start> 1 3 100 -em <GF_end> 3 1 0 -em <GF_end> 1 3 0
 -ej 2 2 1 -ej <t_b> 3 1 -ej 2 5 4 -ej 4 6 4 -ej 2 8 7 -ej 4 9 7
 -ej 8 4 1 -ej 16 7 1
~~~

In these command lines, the parameter *rho* corresponds to the population-scaled recombination rate *ρ*, and `GF start` and `GF end` define the beginning and the end of the pulse of migration in unit of 4*N*_*e*_ generations. We used two values of *ρ*, 1.5 and 15, together with a non-recombining control (*ρ* = 0). The two recombination rates correspond to approximately 3 and 30 times the population-scaled rate expected for a 1,000 bp locus under the average human recombination rate, respectively. In the main text and in Table S1, GF timing is expressed relative to *t*_1_, the time of the *A*–*B* split. Because *t*_1_ = 8*N*_*e*_, this corresponds to *t*_1_ = 2 in the 4*N*_*e*_ time units used by `ms` . The three GF timing categories were therefore implemented as follows:

- recent GF, `GF start` = 0 and `GF end` = 0.2, corresponding to [0, 0.1] relative to *t*_1_;
- intermediate GF, `GF start` = 0.9 and `GF end` = 1.1, corresponding to [0.45, 0.55] relative to *t*_1_;
- for an ancestral GF, `GF start` = 1.79 and `GF end` = 1.99, corresponding to [0.895, 0.995] relative to *t*_1_.

The parameter *t b* corresponds to *t*_2_, the time of the (*A, B*)–*C* split. In `ms`, times are expressed in units of 4*N*_*e*_ generations. Thus, the fixed value *t*_1_=8*Ne* used in the main text corresponds to *τ*_1_=2 in `ms` units, and the three values of *t*_2_, 16*N*_*e*_, 13*N*_*e*_, and 10*N*_*e*_, correspond to *τ*_2_=4, 3.25 and 2.5, respectively. The expected probability of incomplete lineage sorting was calculated as 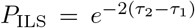, where *τ*_1_ and *τ*_2_ are expressed in `ms` units (Pamilo and Nei, 1988). This gives expected ILS probabilities of approximately 2%, 8% and 37%, respectively:

- *t b* = 4, corresponding to approximately 2% ILS;
- *t b* = 3.25, corresponding to approximately 8% ILS;
- *t b* = 2.5, corresponding to approximately 37%.

Each replicate dataset contained 1,000 loci, and 100 replicate datasets were simulated for each parameter combination. The correspondence between simulation identifiers and parameter combination is provided in Table S2.

### Processing of Simulated Trees

Branch lengths produced by `ms` are expressed in coalescent units. Before running `Aphid`, all branch lengths were converted into substitutions per site by multiplying them by 4.*N*_*e*_*µ*. With *N*_*e*_=10, 000 and *µ*=10^−8^, this scaling factor was 4 *×* 10^−4^.

For non-recombining simulations, the tree produced for each locus was used directly after rescaling. For recombining simulations, `ms` produced one local tree for each non-recombining segment of the locus. These local trees were summarized into a single locus tree using the weighted distance-matrix procedure described in the main text (Fig. 2). The final input file therefore contained one tree per locus, with 1,000 locus trees for each replicate dataset.

### Aphid Analysis and Post-processing

Each replicate dataset was analysed with `Aphid` using the following command:

~~~
Aphid <Gene-Trees_File> <Taxon_File> <Config_File> <Output_File> >>
<stdout>
~~~

The `Gene-Tree File` is a tab-separated file containing one tree per locus, the locus length and a unique locus identifier. The `Taxon File` defined species 1, 2 and 3 as the focal triplet, species 4, 5 and 6 as non-focal species, and species 7, 8 and 9as outgroups. The `Config File` contained the filtering criteria, likelihood settings, and output options used by `Aphid` . The standard output was also redirected into a comma-separated file to obtain dataset-level estimates of the relative contribution of gene flow and incomplete lineage sorting, as well as topological imbalance.

The `Aphid` outputs were then used to extract the proportion of loci inferred to be affected by gene flow, the relative timing of gene flow, and the direction of gene flow, as described in the main text.

`Taxon file` used by Aphid:

~~~
triplet: 1, 2, 3
outgroup: 7, 8, 9
others: 4, 5, 6
~~~

Config file used by Aphid:

~~~
#gene tree filtering
require_triplet_other_monophyly = 1
require_outgroup_monophyly = 0
max_alphai = 10
max_clock_ratio = 2
triplet_clock_thresh = 0.0

#likelihood calculation nb_ILS_times = 1
nb_GF_times = 10
GF_time_coeff = 0.1, 0.2, 0.3, 0.4, 0.5, 0.6, 0.7, 0.8, 0.9, 1.0
min_d = 0.0001
strict_no_event = 0

#likelihood optimisation
max_backward_moves = 100
max_iterations = 100000
convergence = 0.000001

#output
posterior_threshold = 0.95
CI_param = 0
CI_ILS_GF = 0
verbose = 0
~~~

## Supplementary Figures and Tables

**Fig. S1.**
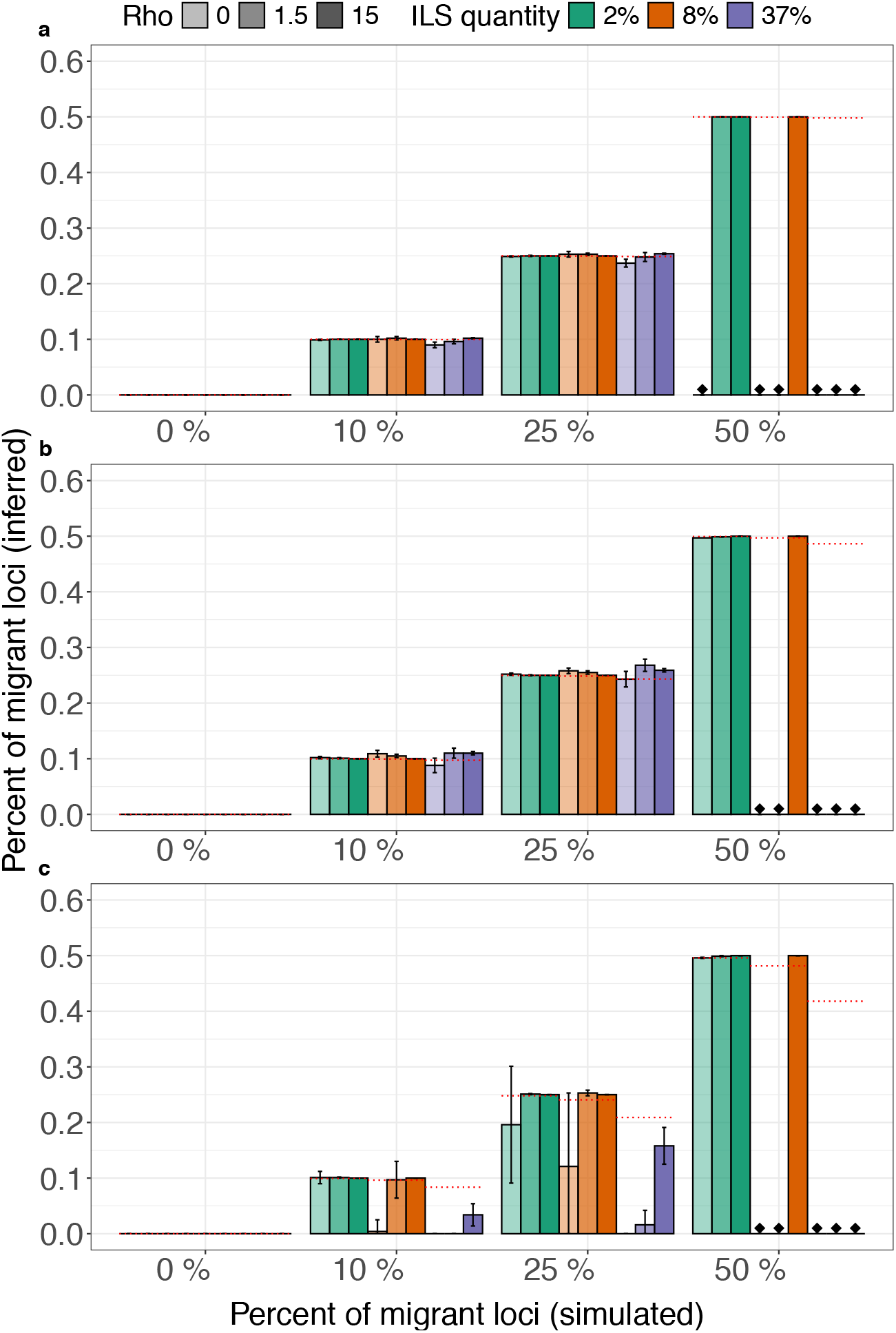
Inferred proportion of loci affected by gene flow (GF) from species A to species C, for recent GF (a), intermediate GF (b) and ancestral GF (c). Bars represent the mean proportion of loci inferred by `Aphid` as affected by GF. Colours indicate the level of ILS: green for 2%, orange for 8%, and blue for 37%. Colours intensity indicates the recombination rate, from the lightest share for *ρ* = 0, to the intermediate shade for *ρ* = 1.5, and the darkest shade for *ρ* = 15. Red dashed lines indicate the expected values under each simulated scenario, accounting for the proportion of loci with discordant topology generated by ILS. Black diamonds indicate parameter combinations for which no replicate dataset satisfied the requirement that at least 50% of analysed trees have the concordant topology.

**Fig. S2.**
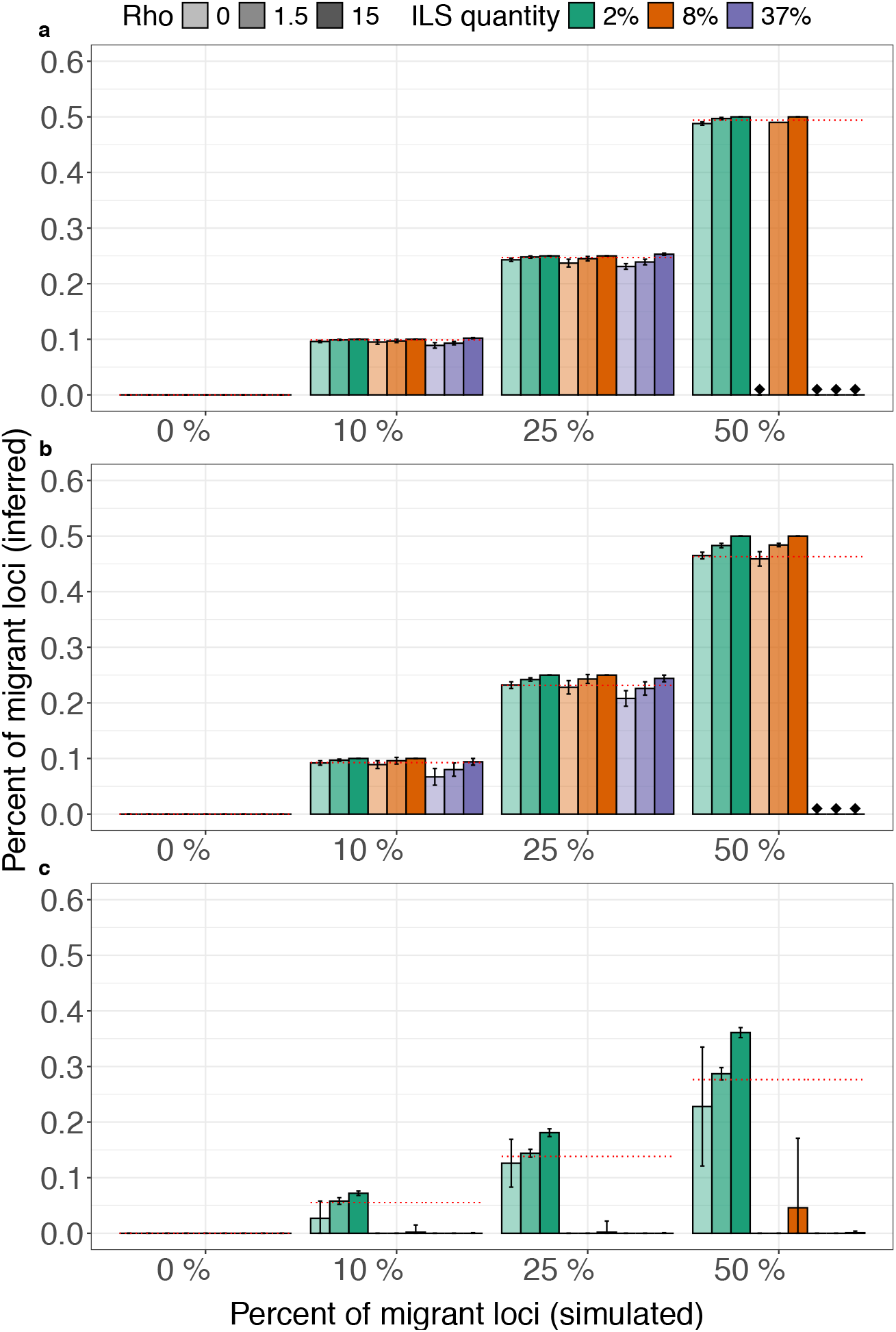
Inferred proportion of loci affected by gene flow (GF) from species *C* to species *A*. Panels, bars, colours, and colour intensities are as in Supplementary Fig. S1.

**Fig. S3.**
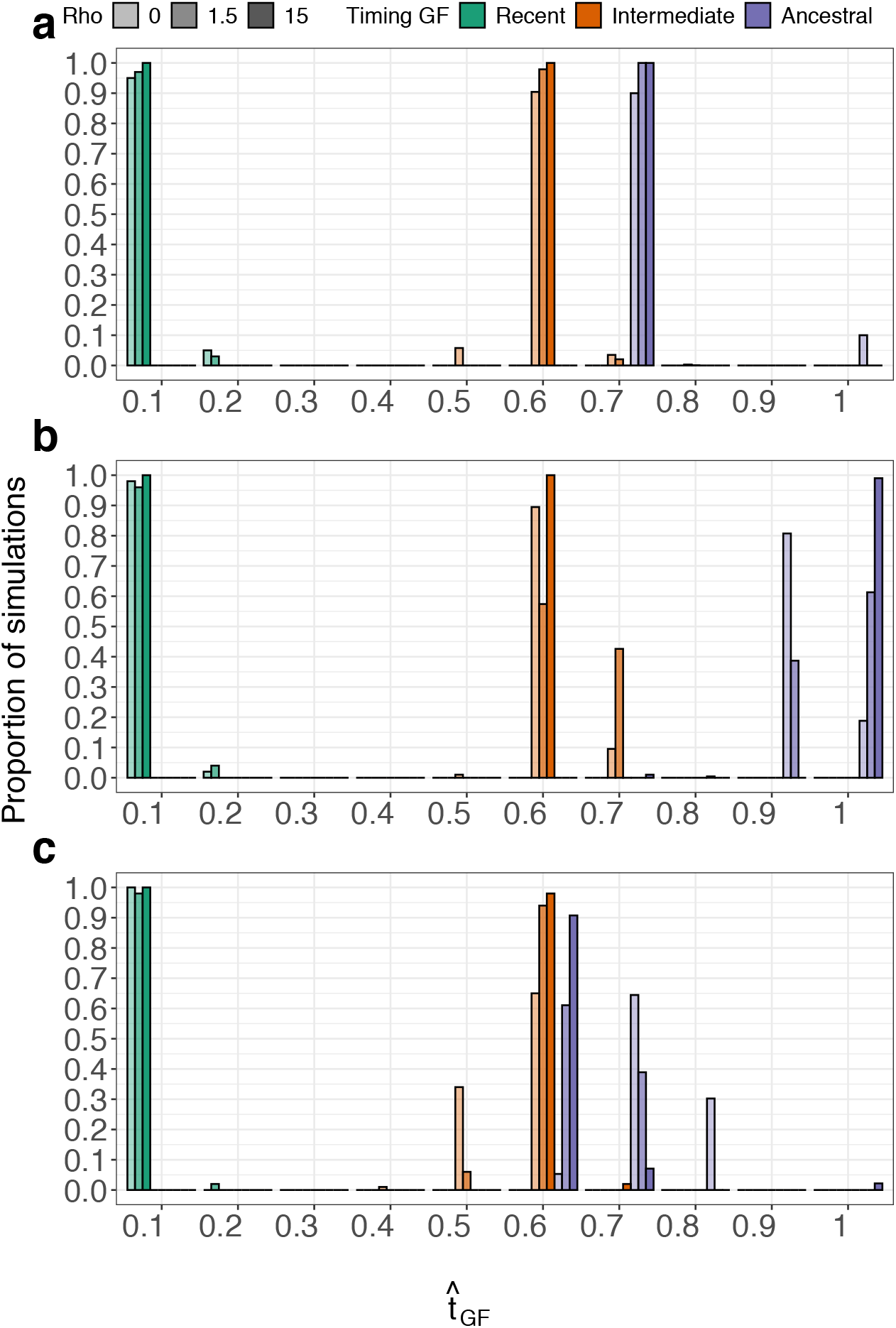
Timing inferred by `Aphid` for simulation in which 25% of loci were affected by GF from species C to species A. Panels correspond to the three levels of ILS: 2% in panel (a), 8% in panel (b), and 37% in panel (c). Bars represent the proportion of replicate datasets assigned to each inferred GF timing category. Colours indicate the simulated timing of GF: green for recent GF, orange for intermediate GF, and blue for ancestral GF. Colour intensity indicates the recombination rate, from the lightest shade for *ρ* = 0, to the intermediate shade for *ρ* = 1.5, and the darkest shade for *ρ* = 15.

**Fig. S4.**
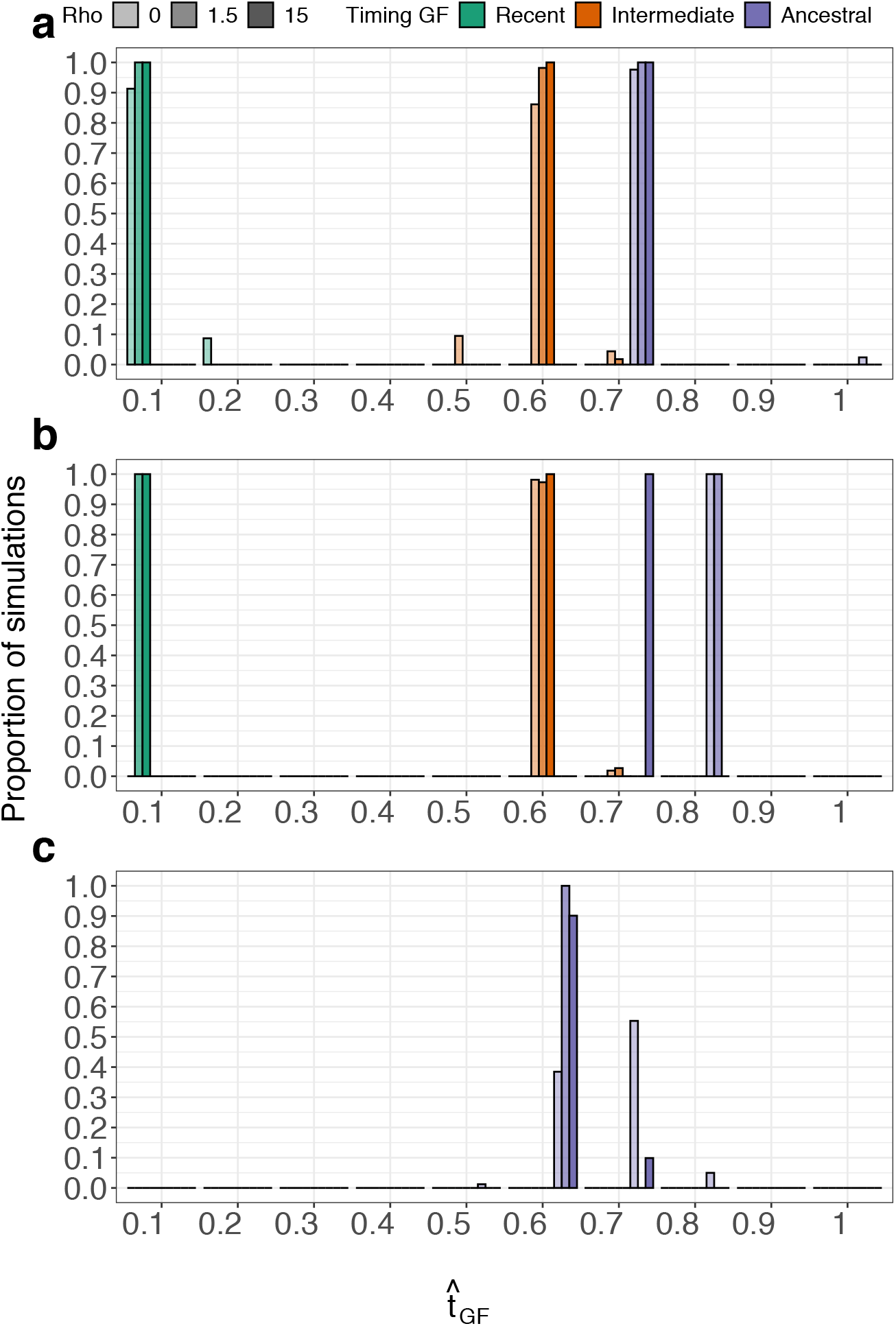
Timing inferred by `Aphid` for simulation in which 50% of loci were affected by GF from species C to species A. Panels, bars, colours, and colour intensities are as in Fig. S3

**Fig. S5.**
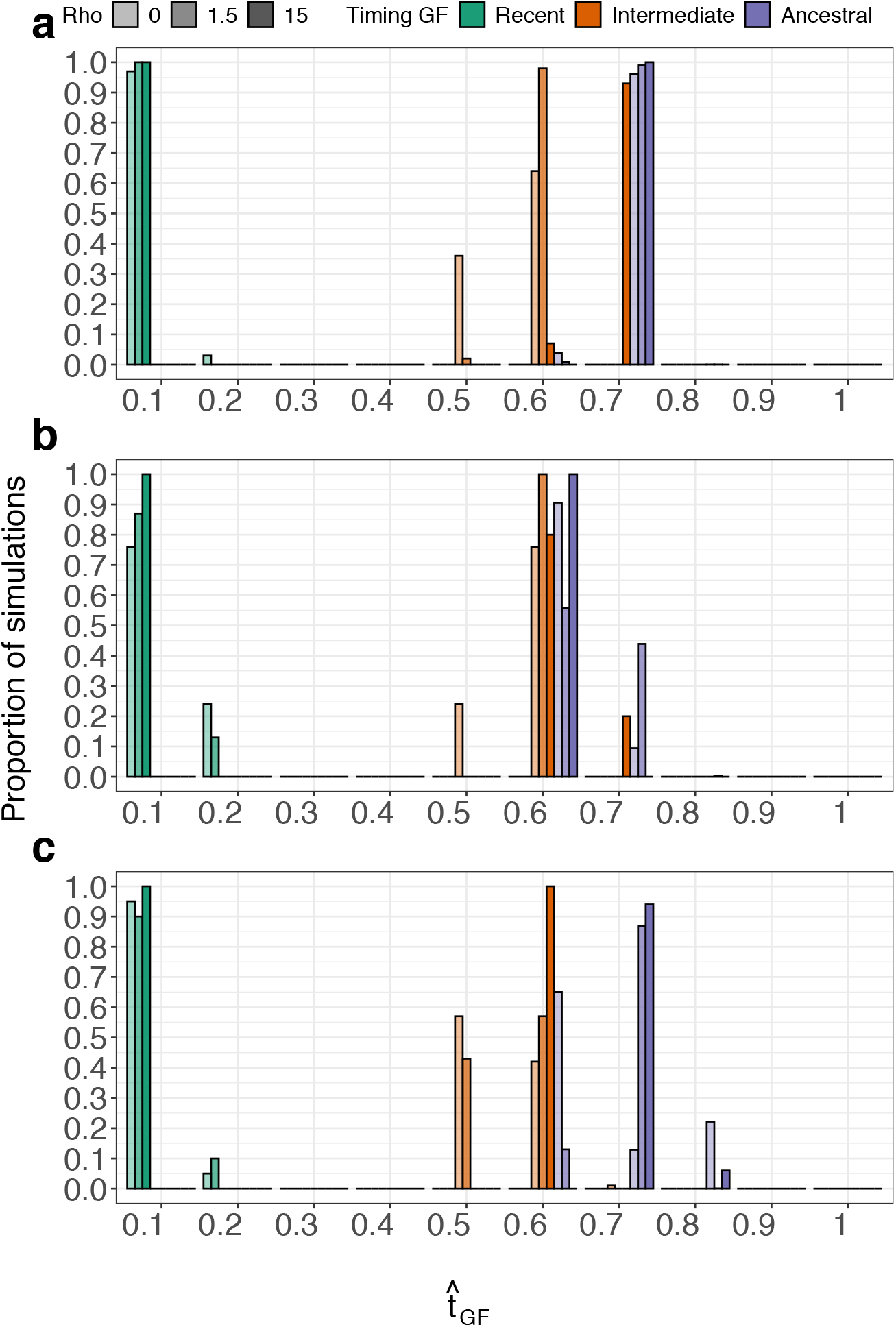
Timing inferred by `Aphid` for simulation in which 25% of loci were affected by GF from species A to species C. Panels, bars, colours, and colour intensities are as in Fig. S3

**Fig. S6.**
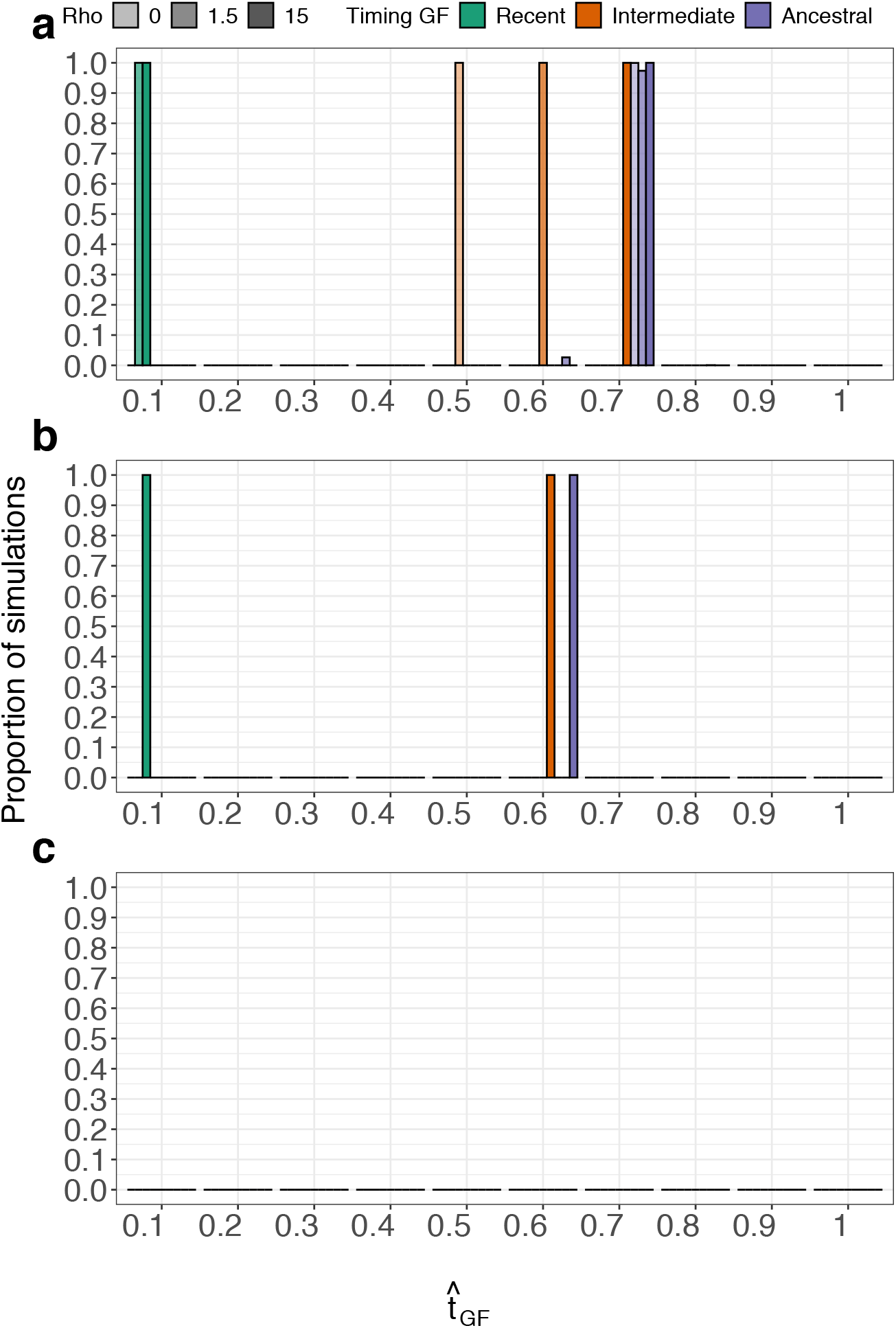
Timing inferred by `Aphid` for simulation in which 50% of loci were affected by GF from species A to species C. Panels, bars, colours, and colour intensities are as in Fig. S3

**Table S2:** Simulation identifiers and associated parameter combinations. Each row corresponds to an identifier of the expected ILS level, GF timing window, backward-in-time direction, proportion of loci affected by GF, and recombination treatment. Species 1 and 3 correspond to species *A* and *C*. Rows marked “Yes” were simulated separately with *ρ*=1.5 and *ρ*=15.

| ID | % ILS | GF model | GF window (relative to $t_1$) | Direction (backward) | % GF | Recombination |
| --- | --- | --- | --- | --- | --- | --- |
| 5.1.b | 2% | None | — | — | 0% | No |
| 5.2.b | 2% | Recent pulse | [0, 0.1] | 3 → 1 | 10% | No |
| 5.3.b | 2% | Intermediate pulse | [0.45, 0.55] | 3 → 1 | 10% | No |
| 5.4.b | 2% | Ancestral pulse | [0.895, 0.995] | 3 → 1 | 10% | No |
| 5.1.d | 2% | None | — | — | 0% | No |
| 5.2.d | 2% | Recent pulse | [0, 0.1] | 3 → 1 | 50% | No |
| 5.3.d | 2% | Intermediate pulse | [0.45, 0.55] | 3 → 1 | 50% | No |
| 5.4.d | 2% | Ancestral pulse | [0.895, 0.995] | 3 → 1 | 50% | No |
| 5.1.e | 2% | None | — | — | 0% | No |
| 5.2.e | 2% | Recent pulse | [0, 0.1] | 3 → 1 | 25% | No |
| 5.3.e | 2% | Intermediate pulse | [0.45, 0.55] | 3 → 1 | 25% | No |
| 5.4.e | 2% | Ancestral pulse | [0.895, 0.995] | 3 → 1 | 25% | No |
| 6.1.b | 2% | None | — | — | 0% | No |
| 6.2.b | 2% | Recent pulse | [0, 0.1] | 1 → 3 | 10% | No |
| 6.3.b | 2% | Intermediate pulse | [0.45, 0.55] | 1 → 3 | 10% | No |
| 6.4.b | 2% | Ancestral pulse | [0.895, 0.995] | 1 → 3 | 10% | No |
| 6.1.d | 2% | None | — | — | 0% | No |
| 6.2.d | 2% | Recent pulse | [0, 0.1] | 1 → 3 | 50% | No |
| 6.3.d | 2% | Intermediate pulse | [0.45, 0.55] | 1 → 3 | 50% | No |
| 6.4.d | 2% | Ancestral pulse | [0.895, 0.995] | 1 → 3 | 50% | No |
| 6.1.e | 2% | None | — | — | 0% | No |
| 6.2.e | 2% | Recent pulse | [0, 0.1] | 1 → 3 | 25% | No |
| 6.3.e | 2% | Intermediate pulse | [0.45, 0.55] | 1 → 3 | 25% | No |
| 6.4.e | 2% | Ancestral pulse | [0.895, 0.995] | 1 → 3 | 25% | No |
| 7.1.b | 2% | None | — | — | 0% | Yes |
| 7.2.b | 2% | Recent pulse | [0, 0.1] | 3 → 1 | 10% | Yes |
| 7.3.b | 2% | Intermediate pulse | [0.45, 0.55] | 3 → 1 | 10% | Yes |
| 7.4.b | 2% | Ancestral pulse | [0.895, 0.995] | 3 → 1 | 10% | Yes |
| 7.1.d | 2% | None | — | — | 0% | Yes |
| 7.2.d | 2% | Recent pulse | [0, 0.1] | 3 → 1 | 50% | Yes |
| 7.3.d | 2% | Intermediate pulse | [0.45, 0.55] | 3 → 1 | 50% | Yes |
| 7.4.d | 2% | Ancestral pulse | [0.895, 0.995] | 3 → 1 | 50% | Yes |
| 7.1.e | 2% | None | — | — | 0% | Yes |
| 7.2.e | 2% | Recent pulse | [0, 0.1] | 3 → 1 | 25% | Yes |
| 7.3.e | 2% | Intermediate pulse | [0.45, 0.55] | 3 → 1 | 25% | Yes |
| 7.4.e | 2% | Ancestral pulse | [0.895, 0.995] | 3 → 1 | 25% | Yes |

Table S2 – Continued from previous page
| ID | % ILS | GF model | GF window (relative to $t_1$) | Direction (backward) | % GF | Recombination |
| --- | --- | --- | --- | --- | --- | --- |
| 8.1.b | 2% | None | – | – | 0% | Yes |
| 8.2.b | 2% | Recent pulse | [0, 0.1] | 1 → 3 | 10% | Yes |
| 8.3.b | 2% | Intermediate pulse | [0.45, 0.55] | 1 → 3 | 10% | Yes |
| 8.4.b | 2% | Ancestral pulse | [0.895, 0.995] | 1 → 3 | 10% | Yes |
| 8.1.d | 2% | None | – | – | 0% | Yes |
| 8.2.d | 2% | Recent pulse | [0, 0.1] | 1 → 3 | 50% | Yes |
| 8.3.d | 2% | Intermediate pulse | [0.45, 0.55] | 1 → 3 | 50% | Yes |
| 8.4.d | 2% | Ancestral pulse | [0.895, 0.995] | 1 → 3 | 50% | Yes |
| 8.1.e | 2% | None | – | – | 0% | Yes |
| 8.2.e | 2% | Recent pulse | [0, 0.1] | 1 → 3 | 25% | Yes |
| 8.3.e | 2% | Intermediate pulse | [0.45, 0.55] | 1 → 3 | 25% | Yes |
| 8.4.e | 2% | Ancestral pulse | [0.895, 0.995] | 1 → 3 | 25% | Yes |
| 14.5.b | 37% | None | – | – | 0% | No |
| 14.6.b | 37% | Recent pulse | [0, 0.1] | 3 → 1 | 10% | No |
| 14.7.b | 37% | Intermediate pulse | [0.45, 0.55] | 3 → 1 | 10% | No |
| 14.8.b | 37% | Ancestral pulse | [0.895, 0.995] | 3 → 1 | 10% | No |
| 14.5.d | 37% | None | – | – | 0% | No |
| 14.6.d | 37% | Recent pulse | [0, 0.1] | 3 → 1 | 50% | No |
| 14.7.d | 37% | Intermediate pulse | [0.45, 0.55] | 3 → 1 | 50% | No |
| 14.8.d | 37% | Ancestral pulse | [0.895, 0.995] | 3 → 1 | 50% | No |
| 14.5.e | 37% | None | – | – | 0% | No |
| 14.6.e | 37% | Recent pulse | [0, 0.1] | 3 → 1 | 25% | No |
| 14.7.e | 37% | Intermediate pulse | [0.45, 0.55] | 3 → 1 | 25% | No |
| 14.8.e | 37% | Ancestral pulse | [0.895, 0.995] | 3 → 1 | 25% | No |
| 15.5.b | 37% | None | – | – | 0% | No |
| 15.6.b | 37% | Recent pulse | [0, 0.1] | 1 → 3 | 10% | No |
| 15.7.b | 37% | Intermediate pulse | [0.45, 0.55] | 1 → 3 | 10% | No |
| 15.8.b | 37% | Ancestral pulse | [0.895, 0.995] | 1 → 3 | 10% | No |
| 15.5.d | 37% | None | – | – | 0% | No |
| 15.6.d | 37% | Recent pulse | [0, 0.1] | 1 → 3 | 50% | No |
| 15.7.d | 37% | Intermediate pulse | [0.45, 0.55] | 1 → 3 | 50% | No |
| 15.8.d | 37% | Ancestral pulse | [0.895, 0.995] | 1 → 3 | 50% | No |
| 15.5.e | 37% | None | – | – | 0% | No |
| 15.6.e | 37% | Recent pulse | [0, 0.1] | 1 → 3 | 25% | No |
| 15.7.e | 37% | Intermediate pulse | [0.45, 0.55] | 1 → 3 | 25% | No |
| 15.8.e | 37% | Ancestral pulse | [0.895, 0.995] | 1 → 3 | 25% | No |
| 16.5.b | 37% | None | – | – | 0% | Yes |
| 16.6.b | 37% | Recent pulse | [0, 0.1] | 3 → 1 | 10% | Yes |
| 16.7.b | 37% | Intermediate pulse | [0.45, 0.55] | 3 → 1 | 10% | Yes |
| 16.8.b | 37% | Ancestral pulse | [0.895, 0.995] | 3 → 1 | 10% | Yes |
| 16.5.d | 37% | None | – | – | 0% | Yes |

Table S2 – Continued from previous page
| ID | % ILS | GF model | GF window (relative to $t_1$) | Direction (backward) | % GF | Recombination |
| --- | --- | --- | --- | --- | --- | --- |
| 16.6_d | 37% | Recent pulse | [0, 0.1] | 3 → 1 | 50% | Yes |
| 16.7_d | 37% | Intermediate pulse | [0.45, 0.55] | 3 → 1 | 50% | Yes |
| 16.8_d | 37% | Ancestral pulse | [0.895, 0.995] | 3 → 1 | 50% | Yes |
| 16.5_e | 37% | None | – | – | 0% | Yes |
| 16.6_e | 37% | Recent pulse | [0, 0.1] | 3 → 1 | 25% | Yes |
| 16.7_e | 37% | Intermediate pulse | [0.45, 0.55] | 3 → 1 | 25% | Yes |
| 16.8_e | 37% | Ancestral pulse | [0.895, 0.995] | 3 → 1 | 25% | Yes |
| 17.5_b | 37% | None | – | – | 0% | Yes |
| 17.6_b | 37% | Recent pulse | [0, 0.1] | 1 → 3 | 10% | Yes |
| 17.7_b | 37% | Intermediate pulse | [0.45, 0.55] | 1 → 3 | 10% | Yes |
| 17.8_b | 37% | Ancestral pulse | [0.895, 0.995] | 1 → 3 | 10% | Yes |
| 17.5_d | 37% | None | – | – | 0% | Yes |
| 17.6_d | 37% | Recent pulse | [0, 0.1] | 1 → 3 | 50% | Yes |
| 17.7_d | 37% | Intermediate pulse | [0.45, 0.55] | 1 → 3 | 50% | Yes |
| 17.8_d | 37% | Ancestral pulse | [0.895, 0.995] | 1 → 3 | 50% | Yes |
| 17.5_e | 37% | None | – | – | 0% | Yes |
| 17.6_e | 37% | Recent pulse | [0, 0.1] | 1 → 3 | 25% | Yes |
| 17.7_e | 37% | Intermediate pulse | [0.45, 0.55] | 1 → 3 | 25% | Yes |
| 17.8_e | 37% | Ancestral pulse | [0.895, 0.995] | 1 → 3 | 25% | Yes |
| 18.5_b | 8% | None | – | – | 0% | No |
| 18.6_b | 8% | Recent pulse | [0, 0.1] | 3 → 1 | 10% | No |
| 18.7_b | 8% | Intermediate pulse | [0.45, 0.55] | 3 → 1 | 10% | No |
| 18.8_b | 8% | Ancestral pulse | [0.895, 0.995] | 3 → 1 | 10% | No |
| 18.5_d | 8% | None | – | – | 0% | No |
| 18.6_d | 8% | Recent pulse | [0, 0.1] | 3 → 1 | 50% | No |
| 18.7_d | 8% | Intermediate pulse | [0.45, 0.55] | 3 → 1 | 50% | No |
| 18.8_d | 8% | Ancestral pulse | [0.895, 0.995] | 3 → 1 | 50% | No |
| 18.5_e | 8% | None | – | – | 0% | No |
| 18.6_e | 8% | Recent pulse | [0, 0.1] | 3 → 1 | 25% | No |
| 18.7_e | 8% | Intermediate pulse | [0.45, 0.55] | 3 → 1 | 25% | No |
| 18.8_e | 8% | Ancestral pulse | [0.895, 0.995] | 3 → 1 | 25% | No |
| 19.5_b | 8% | None | – | – | 0% | No |
| 19.6_b | 8% | Recent pulse | [0, 0.1] | 1 → 3 | 10% | No |
| 19.7_b | 8% | Intermediate pulse | [0.45, 0.55] | 1 → 3 | 10% | No |
| 19.8_b | 8% | Ancestral pulse | [0.895, 0.995] | 1 → 3 | 10% | No |
| 19.5_d | 8% | None | – | – | 0% | No |
| 19.6_d | 8% | Recent pulse | [0, 0.1] | 1 → 3 | 50% | No |
| 19.7_d | 8% | Intermediate pulse | [0.45, 0.55] | 1 → 3 | 50% | No |
| 19.8_d | 8% | Ancestral pulse | [0.895, 0.995] | 1 → 3 | 50% | No |
| 19.5_e | 8% | None | – | – | 0% | No |
| 19.6_e | 8% | Recent pulse | [0, 0.1] | 1 → 3 | 25% | No |

Table S2 – Continued from previous page
| ID | % ILS | GF model | GF window (relative to $t_1$) | Direction (backward) | % GF | Recombination |
| --- | --- | --- | --- | --- | --- | --- |
| 19.7_e | 8% | Intermediate pulse | [0.45, 0.55] | 1 → 3 | 25% | No |
| 19.8_e | 8% | Ancestral pulse | [0.895, 0.995] | 1 → 3 | 25% | No |
| 20.5_b | 8% | None | – | – | 0% | Yes |
| 20.6_b | 8% | Recent pulse | [0, 0.1] | 3 → 1 | 10% | Yes |
| 20.7_b | 8% | Intermediate pulse | [0.45, 0.55] | 3 → 1 | 10% | Yes |
| 20.8_b | 8% | Ancestral pulse | [0.895, 0.995] | 3 → 1 | 10% | Yes |
| 20.5_d | 8% | None | – | – | 0% | Yes |
| 20.6_d | 8% | Recent pulse | [0, 0.1] | 3 → 1 | 50% | Yes |
| 20.7_d | 8% | Intermediate pulse | [0.45, 0.55] | 3 → 1 | 50% | Yes |
| 20.8_d | 8% | Ancestral pulse | [0.895, 0.995] | 3 → 1 | 50% | Yes |
| 20.5_e | 8% | None | – | – | 0% | Yes |
| 20.6_e | 8% | Recent pulse | [0, 0.1] | 3 → 1 | 25% | Yes |
| 20.7_e | 8% | Intermediate pulse | [0.45, 0.55] | 3 → 1 | 25% | Yes |
| 20.8_e | 8% | Ancestral pulse | [0.895, 0.995] | 3 → 1 | 25% | Yes |
| 21.5_b | 8% | None | – | – | 0% | Yes |
| 21.6_b | 8% | Recent pulse | [0, 0.1] | 1 → 3 | 10% | Yes |
| 21.7_b | 8% | Intermediate pulse | [0.45, 0.55] | 1 → 3 | 10% | Yes |
| 21.8_b | 8% | Ancestral pulse | [0.895, 0.995] | 1 → 3 | 10% | Yes |
| 21.5_d | 8% | None | – | – | 0% | Yes |
| 21.6_d | 8% | Recent pulse | [0, 0.1] | 1 → 3 | 50% | Yes |
| 21.7_d | 8% | Intermediate pulse | [0.45, 0.55] | 1 → 3 | 50% | Yes |
| 21.8_d | 8% | Ancestral pulse | [0.895, 0.995] | 1 → 3 | 50% | Yes |
| 21.5_e | 8% | None | – | – | 0% | Yes |
| 21.6_e | 8% | Recent pulse | [0, 0.1] | 1 → 3 | 25% | Yes |
| 21.7_e | 8% | Intermediate pulse | [0.45, 0.55] | 1 → 3 | 25% | Yes |
| 21.8_e | 8% | Ancestral pulse | [0.895, 0.995] | 1 → 3 | 25% | Yes |

**Table S3.** Genealogical complexity within recombining loci. We report the mean number of non-recombining segments per locus their topological composition at *ρ*=1.5 and *ρ*=15, with and without GF. Counts of discordant local trees are shown for different expected ILS level. The final rows give the proportion of loci containing multiple local trees and multiple topologies.

| | | $\rho = 1.5$ | | $\rho = 15$ | |
| --- | --- | --- | --- | --- | --- |
|  |  | GF | No GF | GF | No GF |
| Mean number of non-recombining segments per locus |  | 87.65 | 89.40 | 596.61 | 604.38 |
| Mean number of concordant local trees per locus |  | 23.45 | 80.09 | 158.59 | 541.56 |
| Mean number of discordant local trees per locus | 2% ILS | 62.92 | 1.12 | 428.46 | 7.57 |
|  | 8% ILS | 63.37 | 4.99 | 432.26 | 33.35 |
|  | 37% ILS | 66.32 | 21.83 | 453.35 | 147.56 |
| Proportion of loci with multiple local trees |  | 1.0 | 1.0 | 1.0 | 1.0 |
| Proportion of loci with multiple topologies |  | 0.20 | 0.23 | 0.51 | 0.63 |

**Table S4:** Proportion of replicate datasets in which at least 50% of locus trees had the concordant topology ((*A, B*), *C*). Simulations not shown met this criterion in all 100 replicates. Dashes indicate combinations that were not simulated.

| Simulation ID | $\rho = 0$ | $\rho = 1.5$ | $\rho = 15$ |
| --- | --- | --- | --- |
| 5_2.d | 0.23 | – | – |
| 5_3.d | 1.00 | – | – |
| 5_4.d | 1.00 | – | – |
| 6_2.d | 0.00 | – | – |
| 6_3.d | 0.01 | – | – |
| 6_4.d | 0.10 | – | – |
| 7_2.d | – | 0.63 | 1.00 |
| 7_3.d | – | 1.00 | 1.00 |
| 7_4.d | – | 1.00 | 1.00 |
| 8_2.d | – | 0.36 | 1.00 |
| 8_3.d | – | 0.43 | 1.00 |
| 8_4.d | – | 0.38 | 1.00 |
| 14_6.d | 0.00 | – | – |
| 14_7.d | 0.00 | – | – |
| 14_8.d | 0.20 | – | – |
| 15_6.d | 0.00 | – | – |
| 15_7.d | 0.00 | – | – |
| 15_8.d | 0.00 | – | – |
| 16_6.d | – | 0.00 | 0.00 |
| 16_7.d | – | 0.00 | 0.00 |
| 16_8.d | – | 0.78 | 1.00 |
| 17_6.d | – | 0.00 | 0.00 |
| 17_7.d | – | 0.00 | 0.00 |
| 17_8.d | – | 0.00 | 0.00 |
| 18_6.d | 0.00 | – | – |
| 18_7.d | 0.10 | – | – |
| 18_8.d | 1.00 | – | – |
| 19_6.d | 0.00 | – | – |
| 19_7.d | 0.00 | – | – |
| 19_8.d | 0.00 | – | – |
| 20_6.d | – | 0.01 | 1.00 |
| 20_7.d | – | 0.32 | 1.00 |
| 20_8.d | – | 1.00 | 1.00 |
| 21_6.d | – | 0.00 | 1.00 |
| 21_7.d | – | 0.00 | 0.99 |
| 21_8.d | – | 0.00 | 0.99 |

**Table S5.** Estimated bias, mean absolute error (MAE), and root mean squared error (RMSE) of the inferred proportion of loci aff by GF. Overall estimates are averaged over ILS level, simulated GF proportion, timing, and direction. Timing-specific estimates averaged over ILS level, simulated GF proportion, and direction.

| GF timing | $\rho$ | Bias | MAE | RMSE |
| --- | --- | --- | --- | --- |
| All | 0 | -0.0459 | 0.0491 | 0.0907 |
|  | 1.5 | -0.0315 | 0.0362 | 0.0777 |
|  | 15 | -0.0245 | 0.0263 | 0.0667 |
| Recent | 0 | -0.0067 | 0.0080 | 0.0103 |
|  | 1.5 | -0.0025 | 0.0041 | 0.0058 |
|  | 15 | 0.0010 | 0.0010 | 0.0020 |
| Intermediate | 0 | -0.0101 | 0.0150 | 0.0208 |
|  | 1.5 | -0.0013 | 0.0091 | 0.0136 |
|  | 15 | 0.0006 | 0.0028 | 0.0055 |
| Ancestral | 0 | -0.1209 | 0.1243 | 0.1554 |
|  | 1.5 | -0.0908 | 0.0953 | 0.1338 |
|  | 15 | -0.0751 | 0.0751 | 0.1153 |

Notes: Bias is defined as the inferred minus the simulated proportion of migrant loci.
Overall contrasts between each recombination treatment and $\rho=0$ were significant for all three error measures ( $P < 0.0001$ ).
Within timing categories, the reduction in squared error was significant only for ancestral GF.
$P$ -values were adjusted for simultaneous comparisons using the multivariate- $t$ method.

**Table S6.** Effects of recombination rate on the accuracy of the inferred proportion of loci affected by GF. The overall effect of *ρ* was tested using additive models. Variation in this effect among simulation conditions was assessed by comparing additive models models including interactions between *ρ* and ILS level, simulated GF proportion, timing, and direction.

| Response | Overall effect of $\rho$ | All interactions | $\rho \times \text{ILS}$ | $\rho \times P_{\text{GF}}$ | $\rho \times \text{timing}$ | $\rho \times \text{direction}$ |
| --- | --- | --- | --- | --- | --- | --- |
| Bias | 182.97 | 33.02 | 19.92 | 15.87 | 61.44 | 19.53 |
| Absolute error | 219.44 | 40.74 | 17.26 | 22.72 | 78.66 | 29.85 |
| Squared error | 76.87 | 40.12 | 12.53 | 34.83 | 69.83 | 41.17 |

Notes: Values are $F$ statistics.
Tests of the overall effect of recombination rate had 2 and 10,791 numerator and residual degrees of freedom, respectively.
The comparison between additive and interaction models had 12 and 10,779 degrees of freedom.
Interactions with ILS and timing had 4 degrees of freedom, whereas interactions with GF proportion and direction had 2 degrees of freedom.
All tests were significant at $P < 10^{-7}$ .

**Table S7.** Estimated bias, mean absolute error (MAE), and root mean squared error (RMSE) of the relative timing of GF inferred Aphid. Overall estimates are averaged over ILS level, simulated GF proportion, timing, and direction. Timing-specific estimates averaged over ILS level, simulated GF proportion, and direction.

| GF timing | $\rho$ | Bias | MAE | RMSE |
| --- | --- | --- | --- | --- |
| All | 0 | -0.0206 | 0.0816 | 0.1182 |
|  | 1.5 | -0.0309 | 0.1065 | 0.1453 |
|  | 15 | -0.0167 | 0.1033 | 0.1375 |
| Recent | 0 | 0.0095 | 0.0095 | 0.0337 |
|  | 1.5 | 0.0074 | 0.0074 | 0.0281 |
|  | 15 | 0.0000 | 0.0000 | 0.0000 |
| Intermediate | 0 | 0.0752 | 0.0857 | 0.1013 |
|  | 1.5 | 0.1012 | 0.1018 | 0.1105 |
|  | 15 | 0.1213 | 0.1213 | 0.1281 |
| Ancestral | 0 | -0.1439 | 0.1468 | 0.1723 |
|  | 1.5 | -0.2013 | 0.2103 | 0.2243 |
|  | 15 | -0.1702 | 0.1868 | 0.1986 |

Notes: Bias is defined as the inferred timing minus the reference timing (0.1, 0.5, or 0.9 for recent, intermediate, and ancestral GF, respectively).
Accuracy measures were calculated only for replicate datasets in which a positive timing estimate was obtained.
Overall estimates were obtained from additive models. Timing-specific estimates were obtained from models including interactions between recombination rate and the other simulation factors.
RMSE was calculated as the square root of the estimated mean squared error.

**Table S8.** Effects of recombination rate on the accuracy of relative GF timing inferred by Aphid. The overall effect of *ρ* was tested using additive models. Variation in this effect among simulation conditions was assessed by comparing additive models with m including interactions between *ρ* and ILS level, simulated GF proportion, timing, and direction.

| Response | Overall effect of $\rho$ | All interactions | $\rho \times \text{ILS}$ | $\rho \times P_{\text{GF}}$ | $\rho \times \text{timing}$ | $\rho \times \text{direction}$ |
| --- | --- | --- | --- | --- | --- | --- |
| Bias | 46.51 | 95.73 | 78.11 | 28.16 | 189.57 | 6.45 |
| Absolute error | 218.40 | 105.11 | 81.39 | 20.65 | 174.68 | 106.41 |
| Squared error | 165.16 | 127.60 | 128.85 | 37.84 | 173.90 | 135.27 |

Notes: Values are $F$ statistics.
Tests of the overall effect of recombination rate had 2 and 10,575 numerator and residual degrees of freedom, respectively.
The comparison between additive and interaction models had 12 and 10,563 degrees of freedom.
Interactions with ILS and timing had 4 degrees of freedom, whereas interactions with GF proportion and direction had 2 degrees of freedom.
All tests were significant at $P < 0.002$ ; all except the $\rho \times \text{direction}$ interaction for bias were significant at $P < 10^{-7}$ .

**Table S9.** Proportion of retained replicate datasets for which Aphid produced a positive estimate of relative GF timing.

| GF timing | $\rho$ | Estimable/total | Proportion estimable |
| --- | --- | --- | --- |
| Recent | 0 | 1200/1200 | 1.000 |
|  | 1.5 | 1200/1200 | 1.000 |
|  | 15 | 1200/1200 | 1.000 |
| Intermediate | 0 | 1200/1200 | 1.000 |
|  | 1.5 | 1200/1200 | 1.000 |
|  | 15 | 1200/1200 | 1.000 |
| Ancestral | 0 | 1075/1200 | 0.896 |
|  | 1.5 | 1198/1200 | 0.998 |
|  | 15 | 1111/1200 | 0.926 |

Notes: A timing estimate was considered estimable when it was defined and strictly greater than zero. All non-estimable timings occurred in simulations of ancestral GF. Accuracy statistics in Table S7 are conditional on timing being estimable.

**Table S10.**
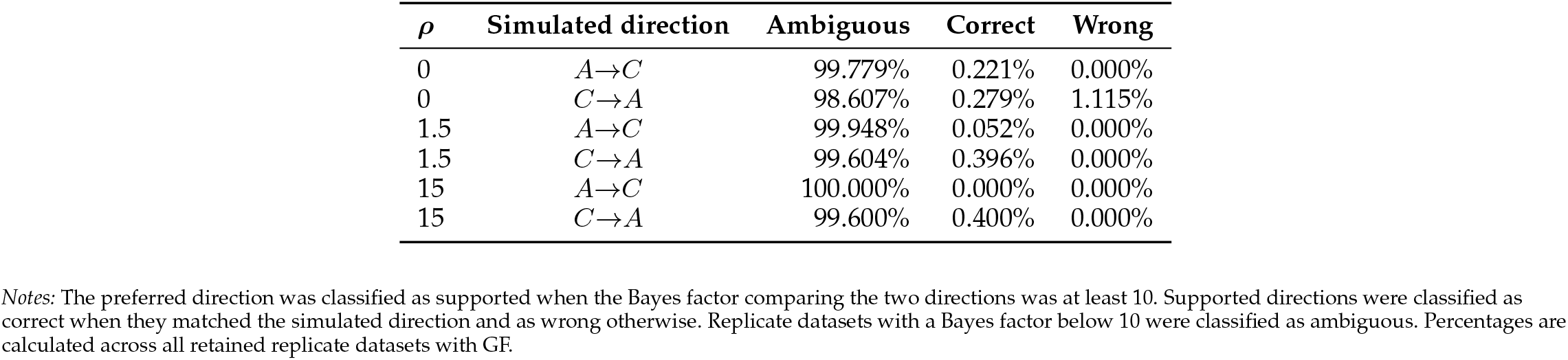
Strength of directional support according to recombination rate.

### Selection of Focal Triplets for the Empirical Cichlid Analysis

The empirical analysis was not intended as an exhaustive scan of all possible species triplets. Instead, we use phylogenetic network inferred by Astudillo-Clavijo et al. (2023) to define a restricted set of biologically motivated parisons. We focused on the reticulation involving *Coptodon* and the Tilapiini–Steatocranini lineage identified in SNaQ analyses (Solís-Lemus and Ané, 2016), and asked whether the corresponding gene-tree discordance conta information about the relative timing of gene flow.

We constructed focal triplets of the form ((*X, Gobiocichla ethelwynnae*), *Coptodon*), where *X* was one of five repr tatives of the lineage potentially involved in gene flow: *Congolapia bilineata, C. crassa, Chilochromis duponti, Steatocr gibbiceps*, or *S. rouxi*. These taxa provide replication at several phylogenetic levels. The two *Congolapia* species test wh the signal is reproducible within this genus, *Chilochromis* tests whether it extends beyond *Congolapia* within Tilapiini the two *Steatocranus* species test whether a similar signal is also present in Steatocranini. A signal restricted to one would therefore be compatible with a more lineage-specific history, whereas a similar signal across several of these would be consistent with a phylogenetically broader event. This taxonomic distribution alone does not date gene timing was inferred from branch lengths with `Aphid` .

Each of the five focal taxa was analysed against four representatives of *Coptodon*: *C. zillii, C. guineensis, C. bakossio* and *C. flavus*. This second level of replication tests whether the inferred signal depends specifically on *C. zillii*, the r sentative used in the original former-tilapiine network, or is recovered with other members of *Coptodon*. Crossing the focal taxa with the four *Coptodon* species resulted in 20 focal triplets (Table S11). The additional control triplets descr in the main Methods were used to provide a reference for the magnitude of 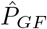, but were not part of this phyloge bracketing strategy.

**Table S11:** Focal triplets used to investigate the relative timing of the *Coptodon*-associated gene-flow signal. All triplets have the reference topology ((*X, Gobiocichla ethelwynnae*), *Coptodon*). The last column indicates the specific purpose of each comparison.

| Focal taxon ($X$) | *Coptodon* | Purpose of the comparison |
| --- | --- | --- |
| *Congolapia bilineata* | *C. zillii* | Direct test of the *Coptodon* signal of GF with a *Congolapia* representative. |
|  | *C. guineensis* | Same comparison with another *Coptodon* (tests whether the signal is specific to *C. zillii*.) |
|  | *C. bakossiorum* | Same comparison with another *Coptodon*. |
|  | *C. flavus* | Same comparison with another *Coptodon*. |
| *Congolapia crassa* | *C. zillii* | Within-*Congolapia* replicate (tests whether the signal is shared by both *Congolapia* species.) |
|  | *C. guineensis* | Same comparison with another *Coptodon*. |
|  | *C. bakossiorum* | Same comparison with another *Coptodon*. |
|  | *C. flavus* | Same comparison with another *Coptodon*. |
| *Chilochromis duponti* | *C. zillii* | Tests whether the signal extends beyond *Congolapia* to another Tilapiini lineage. |
|  | *C. guineensis* | Same comparison with another *Coptodon*. |
|  | *C. bakossiorum* | Same comparison with another *Coptodon*. |
|  | *C. flavus* | Same comparison with another *Coptodon*. |
| *Steatocranus gibbiceps* | *C. zillii* | Tests whether a similar signal extends to Steatocranini. |
|  | *C. guineensis* | Same comparison with another *Coptodon*. |
|  | *C. bakossiorum* | Same comparison with another *Coptodon*. |
|  | *C. flavus* | Same comparison with another *Coptodon*. |

Table S11 – Continued from previous page
| Focal taxon (*X*) | *Coptodon* | Purpose of the comparison |
| --- | --- | --- |
| *Steatocranus rouxi* | *C. zillii* | Independent Steatocranini replicate (tests whether the signal observed with *S. gibbiceps* is species-specific.) |
|  | *C. guineensis* | Same comparison with another *Coptodon*. |
|  | *C. bakossiorum* | Same comparison with another *Coptodon*. |
|  | *C. flavus* | Same comparison with another *Coptodon*. |

For each focal triplet, we considered the discordant topology grouping *X* with *Coptodon*, which is the topology rele to the proposed gene-flow event. Table S12 reports its contributions attributed by `Aphid` to GF and ILS. Because 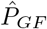 and 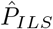 are expressed relative to all retained gene trees, we also report 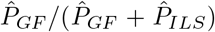, which gives the fra of the focal discordant topology attributed to GF rather than ILS. The relative timing estimate 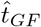 is expressed rel to *τ*_1_, with values approaching zero corresponding to recent GF-associated coalescence and values approaching o coalescence close to the divergence of the two sister lineages (Fig. 6c).

**Table S12.** Detailed Aphid estimates for the 20 focal cichlid triplets. *n* is the number of gene trees retained for each triplet. 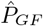 and 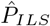 are the dataset-wide contributions of GF and ILS, respectively, for the discordant topology grouping the focal taxon with *Coptodon*. The GF fraction is 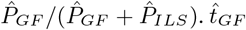 is the GF-associated coalescence time relative to 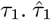 and 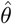 are the corresponding mutation-scaled parameters estimated by `Aphid` .

| Focal taxon (*X*) | *Coptodon* | *n* | $\hat{P}_{GF}$ | $\hat{P}_{ILS}$ | GF fraction within discordance | $\hat{t}_{GF}$ | $\hat{\tau}_1$ | $\hat{\theta}$ |
| --- | --- | --- | --- | --- | --- | --- | --- | --- |
| *Congolapia bilineata* | *C. zillii* | 226 | 0.0327 | 0.1399 | 0.190 | 0.7 | 0.005774 | 0.006693 |
|  | *C. guineensis* | 227 | 0.0427 | 0.1291 | 0.249 | 0.7 | 0.006137 | 0.006398 |
|  | *C. bakossiorum* | 227 | 0.0297 | 0.1333 | 0.182 | 0.8 | 0.006004 | 0.006619 |
|  | *C. flavus* | 227 | 0.0420 | 0.1254 | 0.251 | 0.7 | 0.006247 | 0.006315 |
| *Congolapia crassa* | *C. zillii* | 227 | 0.0302 | 0.1416 | 0.176 | 0.7 | 0.005804 | 0.006746 |
|  | *C. guineensis* | 228 | 0.0378 | 0.1289 | 0.227 | 0.7 | 0.006157 | 0.006476 |
|  | *C. bakossiorum* | 228 | 0.0315 | 0.1352 | 0.189 | 0.7 | 0.006097 | 0.006605 |
|  | *C. flavus* | 228 | 0.0370 | 0.1297 | 0.222 | 0.7 | 0.006221 | 0.006448 |
| *Chilochromis duponti* | *C. zillii* | 226 | 0.0292 | 0.1434 | 0.169 | 0.7 | 0.006855 | 0.006604 |
|  | *C. guineensis* | 227 | 0.0327 | 0.1392 | 0.190 | 0.7 | 0.006906 | 0.006719 |
|  | *C. bakossiorum* | 227 | 0.0283 | 0.1435 | 0.165 | 0.7 | 0.006939 | 0.006692 |
|  | *C. flavus* | 227 | 0.0446 | 0.1448 | 0.235 | 0.7 | 0.006966 | 0.006690 |
| *Steatocranus gibbiceps* | *C. zillii* | 213 | 0.0300 | 0.1531 | 0.164 | 0.7 | 0.008405 | 0.006147 |
|  | *C. guineensis* | 213 | 0.0184 | 0.1600 | 0.103 | 0.7 | 0.008297 | 0.006567 |
|  | *C. bakossiorum* | 213 | 0.0173 | 0.1611 | 0.097 | 0.6 | 0.008343 | 0.006504 |
|  | *C. flavus* | 213 | 0.0182 | 0.1602 | 0.102 | 0.7 | 0.008424 | 0.006405 |
| *Steatocranus rouxi* | *C. zillii* | 210 | 0.0415 | 0.1538 | 0.212 | 0.7 | 0.008930 | 0.005670 |
|  | *C. guineensis* | 209 | 0.0294 | 0.1668 | 0.150 | 0.7 | 0.008833 | 0.006140 |
|  | *C. bakossiorum* | 209 | 0.0265 | 0.1649 | 0.138 | 0.7 | 0.008934 | 0.005998 |
|  | *C. flavus* | 210 | 0.0251 | 0.1607 | 0.135 | 0.7 | 0.008984 | 0.005899 |

